# Reference-free protein sequencing by consensus assembly of redundant de novo peptide reads

**DOI:** 10.64898/2026.08.13.744110

**Authors:** Alfred Nilsson, Emil Sporre, Douwe Schulte, Joost Snijder, Fredrik Edfors, Lukas Käll

## Abstract

Reading a protein’s sequence from tandem mass spectra without a reference is limited by single-spectrum accuracy, most acutely across the hypervariable complementarity-determining regions of antibodies. Broadly specific proteases tile a protein with long, overlapping peptides, so every residue is covered by many independent de novo reads. **borgonovo** assembles their per-step probability profiles into a reference-free per-residue consensus, seeding templates from mass-closure-consistent reads and recruiting the rest by substitution-tolerant alignment and per-column voting. Re-decoding each spectrum with a prior from its consensus position lifts amino acid accuracy on placed spectra from 0.80 to 0.87. On the therapeutic antibody trastuzumab, nine proteases cover its heavy and light chains completely at 0.88 fixed-window identity, and 0.93 on the pruned assembly once local indels are accommodated. Applied unchanged to five secretome proteins and trastuzumab with three proteases, it reaches 0.87 mean fixed-window identity over 82% coverage. borgonovo is open source and works with most de novo sequencers, so redundant digestion turns any of them into a protein sequencer where no reference exists.

## 1 Introduction

Determining a protein’s amino acid sequence directly from the protein, reference-free, without the encoding nucleotide sequence or a database of the expected sequence, is needed whenever that sequence is unknown or unavailable: for engineered and synthetic constructs, for proteins from unsequenced organisms, for forensic and clinical isolates, for biosimilars, and for the variable domains of monoclonal antibodies, which are generated by somatic recombination and so appear in no reference genome.[1] Proteogenomic routes reconstruct such sequences by first sequencing the B-cell repertoire of the same donor and searching against that subject-specific database, which requires matched nucleic acid from the same donor and is not an option for a purified protein of unknown provenance. The need is not confined to sequences that are unknown. Conventional bottom-up analysis identifies peptides against a sequence database and infers the protein by mapping them back onto its entries,[2] so a residue at which the sample’s proteoform departs from an entry is either missed, its peptide matching nothing, or absorbed into the canonical form. Homology-tolerant search relaxes how far a peptide may differ but still requires the database as its alignment target, whereas a sequence read from the spectra requires none. Bottom-up tandem mass spectrometry (MS/MS) is the principal tool, but reading a full-length sequence from it without a reference is hard: a single protease tiles a protein sparsely, and individual de novo spectrum identifications are error-prone. The challenge of reconstructing a complete sequence from noisy de novo reads is shared by any protein one wishes to sequence de novo; antibodies are its most demanding case.

Two recent developments make this newly tractable, and this work combines them. First, broadly specific hyperthermoacidic archaeal (HTA) proteases[3, 4] digest a protein into many long, mutually overlapping peptides, so that every backbone position is sampled by many independent spectra, where a single specific protease provides only a few. They do so in a single 15 min step at one temperature, with no chaotropes, no alkylation and no buffer exchange, so the redundancy costs additional wells rather than additional protocols and the sample preparation is shorter, not longer, than a conventional overnight tryptic digest. Second, deep-learning de novo sequencers, from DeepNovo[5] to Casanovo[6, 7], InstaNovo[8] and Pairwise Attention[9], translate each spectrum directly into a peptide, with no sequence database. Individually these reads remain error-prone, especially in low-complexity or variable regions such as antibody CDRs[10]; but because the broadly specific digest observes each residue in many spectra, the correct residue is almost always present somewhere in the read pile, provided the noisy reads can be placed against one another and aggregated. Assembling that protease-driven redundancy into a finished sequence is the problem addressed here.

Placing short reads against one another, reference-free, is hard when those reads are only *∼*47% accurate per residue (measured on these data below: 0.454 with Casanovo and 0.470 with Pairwise Attention 1.1), and prior approaches fail on it. Classical shotgun protein sequencing by spectral alignment[11] was developed for cleaner input and falters on these noisy reads; its meta-contig consensus[12] and de Bruijn assembly of de novo reads, the route ALPS takes[13], both build contigs from exact or near-exact overlaps that *∼*47%-accurate reads rarely support, so the graph fragments. Cumulative-mass (spectrum-graph) assembly is brittle: one mis-called residue shifts the entire downstream mass ladder out of register, and greedy overlap–layout–consensus collapses when a single spurious transitive overlap fuses unrelated regions. What works instead is to align noisy reads to a clean consensus template and vote per column, but no such template exists a priori in the reference-free setting.

The established way around this problem is to supply the template externally. Template-based assemblers such as Stitch map de novo reads onto user-defined germline (IMGT[14]) V/J/C segments and read off a per-column consensus, reconstructing antibody chains to high accuracy and profiling polyclonal repertoires.[15] But this is contingent on a close reference existing, the right species and alleles of an antibody germline, which is unavailable for an engineered or unknown-organism protein and absent altogether outside the immunoglobulin framework; it also leaves the most template-divergent regions, such as the CDRH3, the hardest to recover. borgonovo instead derives the template from the reads themselves, using no germline or reference sequence at any stage, so a single procedure applies to any protein, whether or not it is an antibody.

Database-independent full-length sequencing has been approached by scaffolding overlapping de novo contigs across multiple unspecific proteases without a germline template.[16, 17] Most recently, Xin et al. assembled polyclonal antibody sequences from overlapping bottom-up de novo reads via a mass-block-tolerant overlap graph, integrating middle-down and intact-mass data to validate and correct the output.[18] borgonovo shares the multi-protease-redundancy thesis but, rather than stitching contigs by sequence or mass-block overlap, distils a single per- column consensus from the reads’ full softmax distributions and feeds that consensus back into the decoder, so the redundancy is exploited both to build the template and to repair the individual reads against it, all from bottom-up data alone, without middle-down or intact-mass integration. Here we present **borgonovo**, which bootstraps the template from the cleanest reads and then recruits the rest. The assembled consensus serves two purposes: it is itself the reconstructed sequence, and it provides a per-position prior that, blended into the de novo model at a second decoding pass, repairs individual spectrum identifications. We demonstrate it reference-free on a panel of non-antibody proteins digested with three broadly specific proteases and on the therapeutic antibody trastuzumab, showing that a single template-free procedure spans both. Because no germline, IMGT or database enters at any stage, nothing in the procedure depends on the sequence being known.

## 2 Results

**borgonovo** is the open-source package implementing the method. It does not sequence spectra itself: it wraps an existing de novo peptide sequencer, treated as an interchangeable backend, and adds the assembly and re-decoding logic around it. Throughout the results the backend is Pairwise Attention 1.1, a model released with this work, with the widely used Casanovo serving later as a substitution control that separates what the backend contributes from what the assembler does. Figure 1 summarizes the three stages the results below follow: first-pass profiling of every spectrum into a per-step probability profile, reference-free assembly of those profiles into per-residue consensus templates, and re-decoding of each spectrum under a prior drawn from the consensus at its placed position. The Methods give each stage in full.

**Figure 1:**
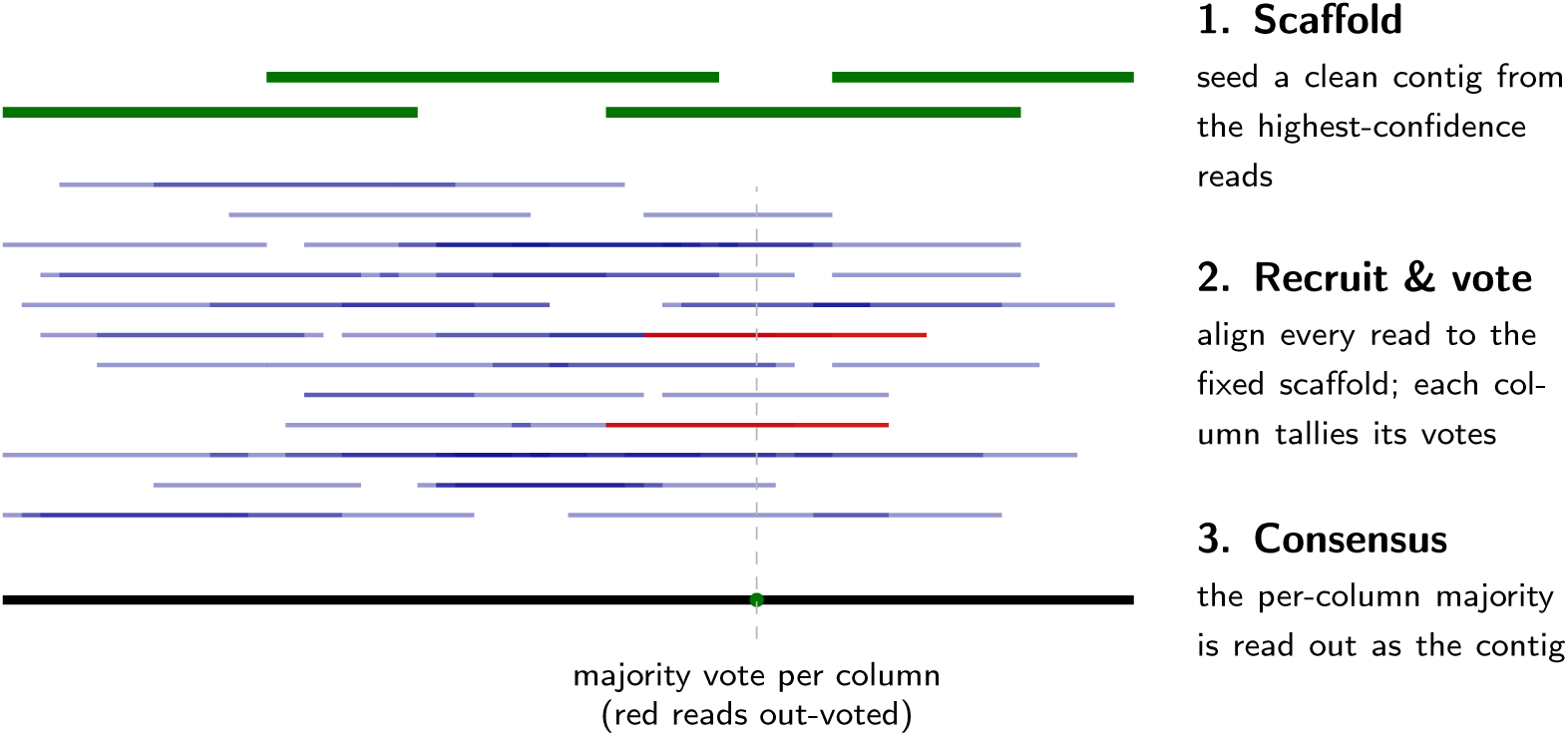
Reference-free assembty by scaffotd-and-recruit. (illustrative; reads are schematic, not a specific protein). A broadly specific digest samples every backbone position with many overlapping de novo reads. **(1)** The cleanest reads, those passing mass closure with the highest confidence, seed a high-confidence consensus *scaffold* (green). **(2)** Every read is then aligned to the fixed scaffold and votes into its columns (substitution-tolerant, no end-extension); at any column a minority miscall (red) is out-voted by the correct majority. **(3)** Each column’s majority call is read out as the consensus contig (black). The same per-column vote distributions double as the prior that biases the second-pass re-decoding of individual spectra.

### 2.1 Reference-free reconstruction of the Fab

Every spectrum is first interpreted on its own into a read, and it is these reads, not the spectra, that the assembler consumes. From the pooled nine-protease trastuzumab data, borgonovo assembles 44 consensus templates in *∼*20 s, covering 100.0% of the heavy chain and 100.0% of the light chain at *≥*50% local identity. Local identity here is a purity filter on each contig rather than a statement about the quality of the coverage: a contig is placed at its single best ungapped offset in a chain, its identity is taken over its whole length so that any overhang counts against it, and only contigs matching at *≥*50% are allowed to contribute coverage at all. Coverage is then the fraction of chain positions lying under at least one such contig, and the per-residue accuracy of those positions is reported separately below. The templates reconstruct the framework regions and the CDRs: for example the heavy-chain CDR1– CDR2 stretch (GFN[L/I]KDTYLHWVRQAPGKGLEWVARIYPTNGYTRYADSVKG) and the light-chain CDR3 (QQHYTTPPTFGQGTKVELK) are recovered at 0.96–0.98 identity. This is achieved without any reference: the only role of mass closure is to choose which reads seed the scaffold. Figure 2 shows the reconstruction along both chains: the reference bar is coloured per residue as correctly identified, incorrectly identified, or not covered, with the matching reads piled above it (shaded by local depth). The error positions cluster in the hypervariable CDRs and coincide with the shallower points of the read pileup, while the constant framework is reconstructed almost perfectly; with nine proteases both chains are covered end to end, so every residue is called.

**Figure 2:**
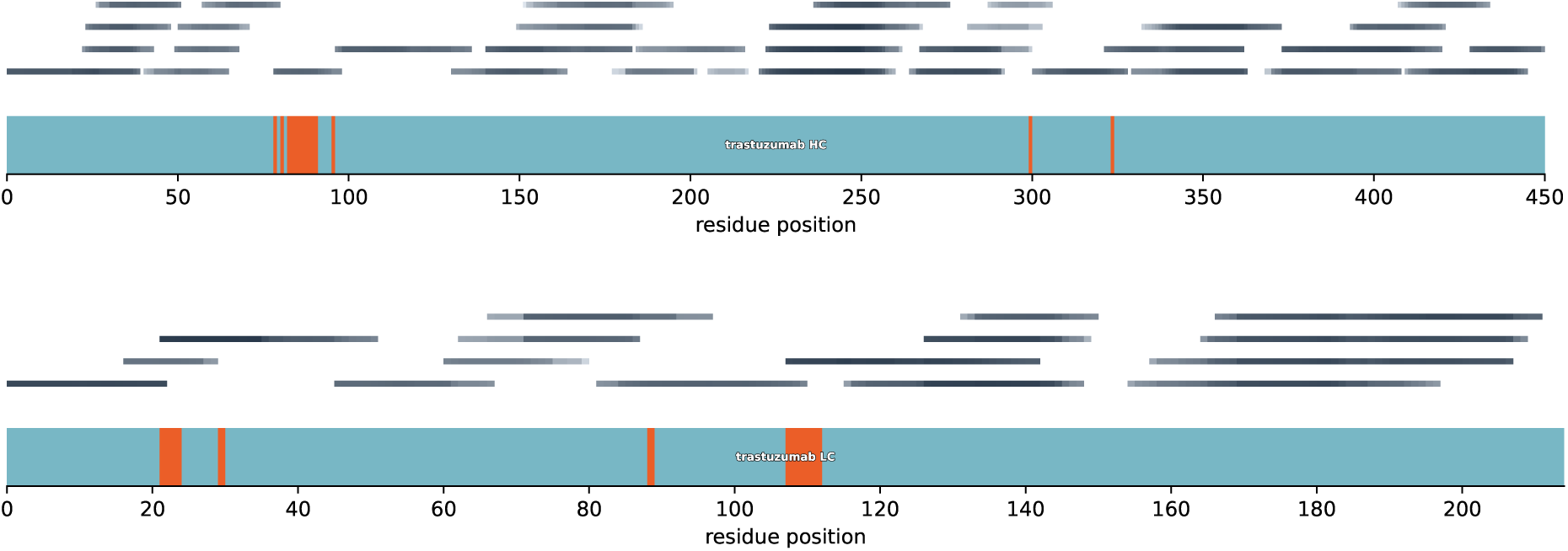
Reference-free reconstruction of the trastuzumab heavy (top) and light (bottom) chains from the pooled *nine-protease* data, scored against the known sequence (used only to colour the figure). Trastuzumab also appears in Figure 3 and Table 3 under a three-protease HTA digest, which is a different and much shallower dataset; the two are not directly comparable. The horizontal bar is the chain, coloured per residue as correctly identified, incorrectly identified, or not covered by any placed read; thin lines above are the matching reads (sub-spans collapsed into their longest containing read, each shaded darker where more reads were collapsed under it). The heavy chain is fully covered with 97% of covered residues correct (17 314 placed reads); the light chain is likewise fully covered with 95% correct (11 286 reads). Errors concentrate in the CDRs, where coverage is shallowest, while the constant framework is near-perfect.

**Table 1:** Reference-free assembly accuracy of trastuzumab (length-weighted over 44 templates; –prune-frac 0.4). Fixed-window: exact residue identity, contiguous, no indels. Pruned: lowdepth columns dropped. Fair: banded alignment absorbing local indels. Mass-aware: additionally credits isobaric blocks. The last two rows re-score the pruned assembly under a more forgiving alignment; coverage is a property of the assembly rather than of the scoring, so it is quoted once per ass<u>embly and left blank (—) where it is unchanged from t</u>he row above.

| scoring | identity | HC cov. | LC cov. |
| --- | --- | --- | --- |
| fixed-window | 0.88 | 100.0% | 100.0% |
| fixed-window, pruned | 0.89 | 94.9% | 98.6% |
| fair (indel-absorbing) | 0.93 | — | — |
| mass-aware | 0.94 | — | — |

**Table 2:** Effect of the consensus prior on per-spectrum amino acid accuracy (tryptic trastuzumab; *α* = 0.5 vs. *α* = 0 co<u>ntrol, identical greedy decoder).</u>

| spectra | $\alpha = 0$ | $\alpha = 0.5$ | $\Delta$ |
| --- | --- | --- | --- |
| all (n=10221) | 0.525 | 0.554 | +0.029 |
| placed (n=4095) | 0.802 | 0.874 | +0.073 |
| prior-changed (n=2351) | 0.715 | 0.841 | +0.126 |
| % AA $\geq 0.9$ (all) | 22.9 | 29.8 | — |

**Table 3:** Reference-free reconstruction of six known proteins from three pooled proteases (krakatoa / vesuvius / HTA3), scored per contig against canonical UniProt. Fixed-window: exact contiguous identity; fair: banded, indel-absorbing; mass-aware: additionally crediting isobaric blocks; coverage: reference fraction under an assigned contig at *≥*0.5 identity. (The per-residue colours in Figure 3 use the deepest covering consensus and so run higher than the per-contig mean.)

| protein | residues | coverage | fixed | fair | mass |
| --- | --- | --- | --- | --- | --- |
| apolipoprotein A-IV (APOA4) | 396 | 96% | 0.85 | 0.93 | 0.95 |
| paraoxonase-1 (PON1) | 355 | 84% | 0.87 | 0.92 | 0.93 |
| factor XI (F11) | 625 | 89% | 0.88 | 0.92 | 0.93 |
| inhibin βC (INHBC) | 352 | 75% | 0.80 | 0.89 | 0.91 |
| C1 inhibitor (SERPING1) | 500 | 70% | 0.90 | 0.96 | 0.97 |
| trastuzumab (HC+LC) | 664 | 80% | 0.94 | 0.97 | 0.98 |
| mean |  | 82% | 0.87 | 0.93 | 0.95 |

Table 1 summarizes accuracy under three scorings. A naive fixed-window residue identity reads 0.88; pruning low-depth (misframed) template columns raises it to 0.89, at a cost in coverage (both chains are already complete before pruning; the heavy chain falls to 94.9% and the light chain to 98.6%). Separately, switching the recruiter itself to the isobaric-tolerant aligner (the placement use of the same equal-mass move; Methods) yields a better assembly: it lifts the pruned fixed-window identity from 0.89 to 0.93 and the corresponding mass-aware identity from 0.94 to 0.96, by keeping reads that tokenize a residue differently from the template in frame. Both recruiters cover the heavy chain completely before pruning, so the gain is in identity rather than extent. We leave it off by default for its added alignment cost. A *fair* banded alignment that absorbs local indels puts the true residue identity at 0.93: the fixed-window number under- reports by *∼*0.05 purely because it cannot accommodate a single insertion or deletion. Crediting mass-isobaric tokenizations adds only a further *∼*0.01, so isobaric ambiguity (Q*≡*GA, GG*≡*N) is a minor contributor. The residual, *∼*6%, is correlated model error (positions where most reads agree on a wrong call, which consensus therefore cannot fix), and it is spread thinly rather than concentrated in one systematic confusion: over the 664 covered positions only 24 are miscalled, and no single substitution accounts for more than two of them. Cysteine is no longer the outlier it is for a backend that cannot emit the unmodified residue: it is called correctly at 13 of 16 covered positions, and there are no carbamidomethyl-cysteine read as alanine, which is the most frequent single mismatch under Casanovo. For context, we first tried to assemble these reads the conventional way, by overlapping them directly with one another instead of against a template. Several variants of that idea all failed, reaching only 0.14–0.41 identity, below the accuracy of the reads going in. A better backend does not rescue them: with Pairwise Attention 1.1 reads the same approach gets worse as more reads are added, 0.16, 0.16 and 0.11 at 100, 200 and 400 reads, because each extra read creates more chances for a spurious overlap to fuse unrelated regions. The bootstrap is therefore not an optimisation but the thing that makes reference-free assembly work at this read accuracy.

### 2.2 The consensus prior improves per-spectrum decoding

To isolate the effect of the consensus prior, we re-decoded the tryptic trastuzumab spectra twice through the identical greedy code path, at *α* = 0 (no prior) and *α* = 0.5, and scored each PSM against the reference (Table 2). Of 10221 decoded spectra, 4095 were matched to a template placement; the prior changed the output of 2351. On those 2351 spectra mean amino acid accuracy rose from 0.72 to 0.84 (+0.126), with 1638 improved versus 162 worsened (an *∼*10:1 ratio), and the fraction of near-perfect identifications (*≥*0.9) rose from 28.0% to 57.8% (over all 10221 spectra the same fraction rises from 22.9% to 29.8%, the value tabulated in Table 2). Averaged over all spectra the gain is +0.03 (diluted by the unplaced majority); a sign test on the 1638:162 improved:worsened split is highly significant (*p ≪* 10*^−^*^300^). The consensus thus repairs individual decodes where it has support, without degrading the rest.

Two caveats could bound this claim. First, the tryptic spectra above also contributed (as one of nine proteases) to the consensus, so the prior was not strictly independent of the spectra it re-decodes, though the per-spectrum leakage is small (one vote in *∼*240-deep columns). Second, the prior-changed subset is, by construction, the spectra whose first pass disagreed with the consensus, which biases toward the consensus being right. To remove both, we built a consensus from the *eight* non-tryptic proteases and placed the tryptic reads against that frozen consensus *without letting them vote*, so no tryptic spectrum contributed to the column that primes it. Re-decoding the tryptic spectra against this fully independent prior reproduces the gain: across all 4702 placed spectra mean accuracy rose 0.74 *→* 0.83 (+0.084), and on the 3112 prior-changed spectra 0.65 *→* 0.78 (+0.126), with 2101 improved versus 522 worsened. The mean gain on the prior-changed subset is the same as in-sample to three decimals, and on all placed spectra it is larger; the improvement ratio is lower (*∼*4:1 against *∼*10:1), so a wholly independent prior helps as much on average but misfires more often. The gain is thus not an artifact of a spectrum’s own read leaking into its prior; and because the all-placed figure is not conditioned on the prior having changed the output, it also escapes the selection bias of the changed-only subset. The improvement comes from the redundancy across proteases, not from re-using a spectrum’s own read.

### 2.3 Iterating the consensus back does not improve the reconstruction

Because the consensus prior measurably improves individual decodes, an obvious extension is to feed those refined decodes back: re-decode every spectrum with the prior, re-assemble the refined reads, and iterate. We tested a single such iteration on trastuzumab. Re-decoding each spectrum along the prior-steered path, we re-assembled either the model’s own per-step softmax along that path or the blended softmax used to steer it, swept the blend weight *α*, and additionally tried a mass-closure gate that accepts a refined read only if its decoded ladder still explains the precursor mass.

No variant, at any *α*, improves on the single-pass consensus (length-weighted fair identity 0.923); the best result, a gentle *α* = 0.1 with the mass-closure gate, merely breaks even (0.924). Without the gate the reconstruction degrades as the prior strengthens and collapses once the prior begins to override the spectrum rather than merely bias it (fair identity *∼*0.70 at *α* = 0.75, whether the model or the blended softmax is re-assembled). The mass-closure gate prevents this collapse (it discards any refined read whose mass ladder no longer fits the precursor, so the result never falls below baseline at any *α*), but it does not lift it above baseline either. The reason is structural: a single consensus pass already averages out the uncorrelated per-spectrum errors, so re-decoding the reads merely re-cleans noise the consensus has already absorbed, while the correlated errors survive the loop untouched; and once a read has been pulled toward the consensus, re-voting it double-counts that consensus rather than contributing independent evidence. The prior’s value therefore lies in repairing individual spectrum identifications, not in refining the consensus: the consensus is best read out once, before re-decoding, and re-decoding never feeds back into the assembled contigs.

### 2.4 Generalization to non-antibody proteins with three proteases

To test the method beyond a single antibody and a single digest, we applied borgonovo unchanged to six known proteins: five human plasma/secretome proteins (apolipoprotein A-IV, paraoxonase-1, the C1 inhibitor SERPING1, coagulation factor XI and inhibin *β*C) and the trastuzumab antibody, each digested separately by three broadly specific proteases (krakatoa, vesuvius and HTA3) and the digests pooled. The per-residue accuracy transfers intact: a mean 0.87 fixed-window, 0.93 fair (indel-absorbing) and 0.95 mass-aware identity over the covered regions (Table 3), the same as on trastuzumab, on proteins of 350–660 residues. Coverage is high where peptide yield is high (apolipoprotein A-IV and paraoxonase-1 at 85–90%) and protein-limited elsewhere, as before.

Running three proteases separately makes the source of that coverage explicit (Figure 3). Collapsing each protease’s reads on its own shows krakatoa contributing the densest peptide tiling, vesuvius a largely complementary one, and HTA3 a sparser overlay; the pooled consensus covers more of each protein than any single protease, and its errors fall where all three are shallow. This is the multi-protease redundancy the method relies on, now shown on non-antibody proteins. The particular choice of the three proteases is what makes that redundancy cheap to obtain.

**Figure 3:**
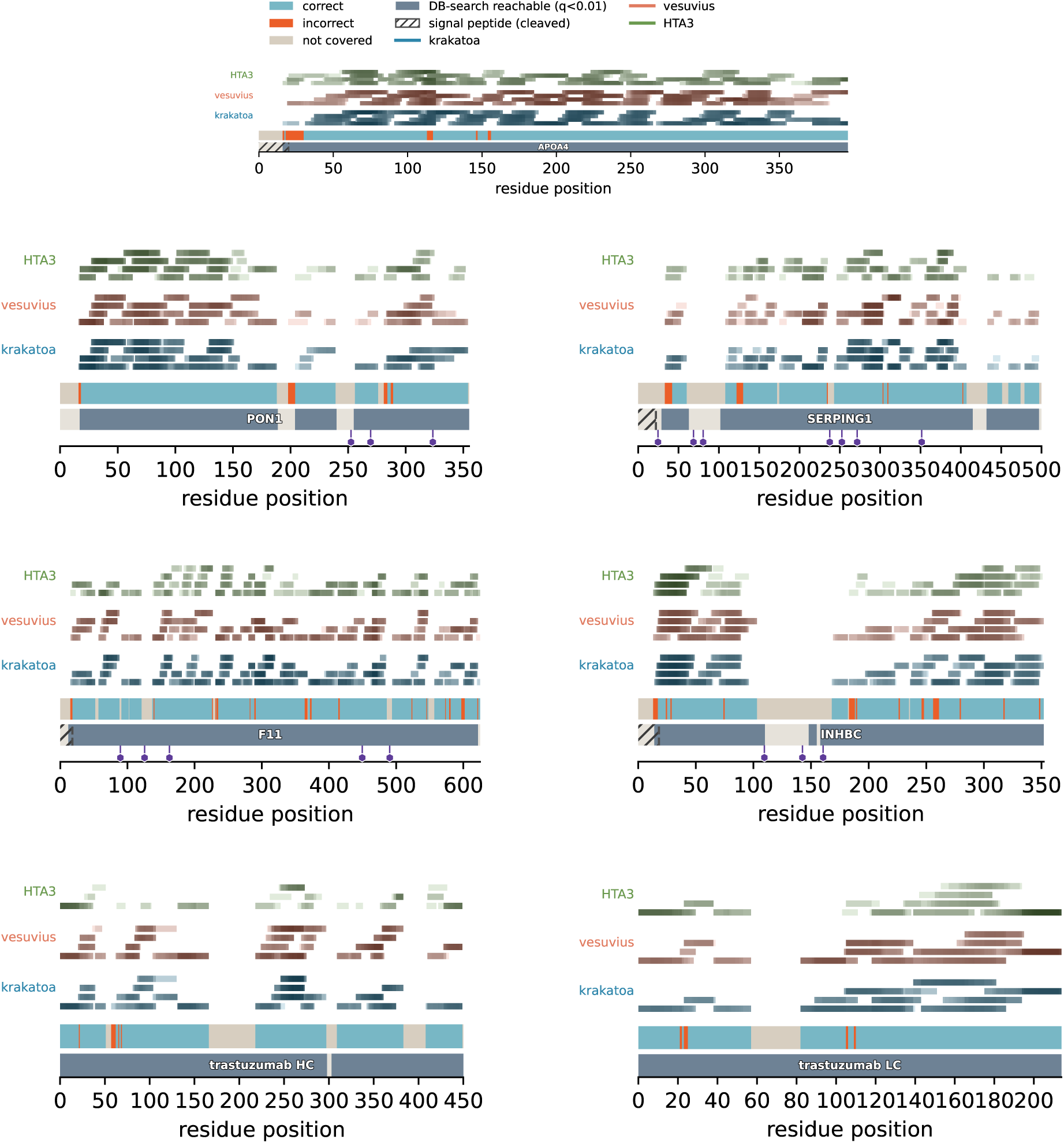
Per-protease reference-free coverage of five secretome proteins and the trastuzumab antibody. Apolipoprotein A-IV is the top (legend) panel; the remaining panels read down the columns: paraoxonase-1, factor XI, trastuzumab heavy chain (left); SERPING1, inhibin *β*C, trastuzumab light chain (right). The thin bar is the reference (correct / incorrect / not covered); above it the reads are piled up *separately for each protease* (krakatoa, vesuvius, HTA3), pooled in Table 3. The track beneath each bar marks where a confident (*q <* 0.01) database search against the known sequence places a peptide, the sequenceable ceiling, most of which the consensus recovers. Its gaps are explained by the annotations on it: a hatch over a cleaved signal peptide (UniProt, Phobius-confirmed; coverage starts at the mature N-terminus; paraoxonase-1, unmarked, retains its signal as a lipoprotein anchor) and violet lollipops at N-linked glycosylation sites, whose glycan mass-shifts the peptide off a bare-sequence match (11/18 sites in uncovered positions vs. *∼*22% uncovered overall; deglycosylation recovers three of them, Figure 4). With only three proteases the trastuzumab chains are the hardest case (heavy 79%, light 88% covered) though most is database-reachable (nine proteases reach 100/100%, Fig. 2). Only the top panel carries the legend.

Krakatoa, vesuvius and HTA3 all operate under the same hyperthermoacidic conditions, so the three digests are set up in the same buffer, in the same plate, at the same temperature and for the same 15 min, with no buffer exchange or protease-specific optimization in between; running three proteases instead of one costs three wells rather than three protocols. The same conditions also remove the reduction–alkylation step: at pH 2–4 the TCEP-reduced cysteine thiols are protonated rather than present as the thiolate that thiol–disulfide exchange requires, so the disulfides do not re-form over the 15 min digest, and TCEP itself stays effective at that pH where dithiothreitol would not. No iodoacetamide or chloroacetamide capping, and no subsequent removal of excess reagent, is therefore needed. Multi-protease coverage is therefore obtained within a sample-preparation protocol that is shorter, not longer, than a conventional single overnight tryptic digest. The one price is the free-thiol cysteine the missing alkylation leaves behind, which the backend model, trained on carbamidomethylated data, systematically miscalls; we return to this in the Discussion.

The remaining sequence-intrinsic gap is glycosylation. An N-linked glycan stays on the peptide and shifts it out of any bare-sequence match, so occupied sequons are systematically under-covered: across the five secretome proteins only 7 of the 18 annotated N-glycosylation sites are assembled from the three proteases, against *∼*22% of residues uncovered overall. To read through them we digested a PNGaseF-deglycosylated aliquot of each protein with krakatoa (PNGaseF releases the glycan and converts the sequon asparagine to aspartate) and assembled it together with the three glycosylated digests (Figure 4). This doubles the covered sequons to 10 of 18, and the recovery is specific to the *occupied* sites: every sequon gained reads the deamidated aspartate, whereas the sites already covered by the glycosylated digests read unmodified asparagine, their non-glycosylated peptidoforms. The gain is a deglycosylation signal, not a decoding artefact: across all covered asparagines the aspartate call is confined to the sequons (an *∼*0.8 rate at N-x-S/T versus none elsewhere). The six sequons still missed lie in regions covered by no digest, so what limits them is the absence of reads rather than a failure to read a deglycosylated site. We cannot conclude that the glycan is irrelevant there: an occupied sequon can itself be why a region yields no usable peptide, through impaired cleavage or suppressed ionisation, in which case the missing coverage is a downstream consequence of glycosylation rather than independent of it.

**Figure 4:**
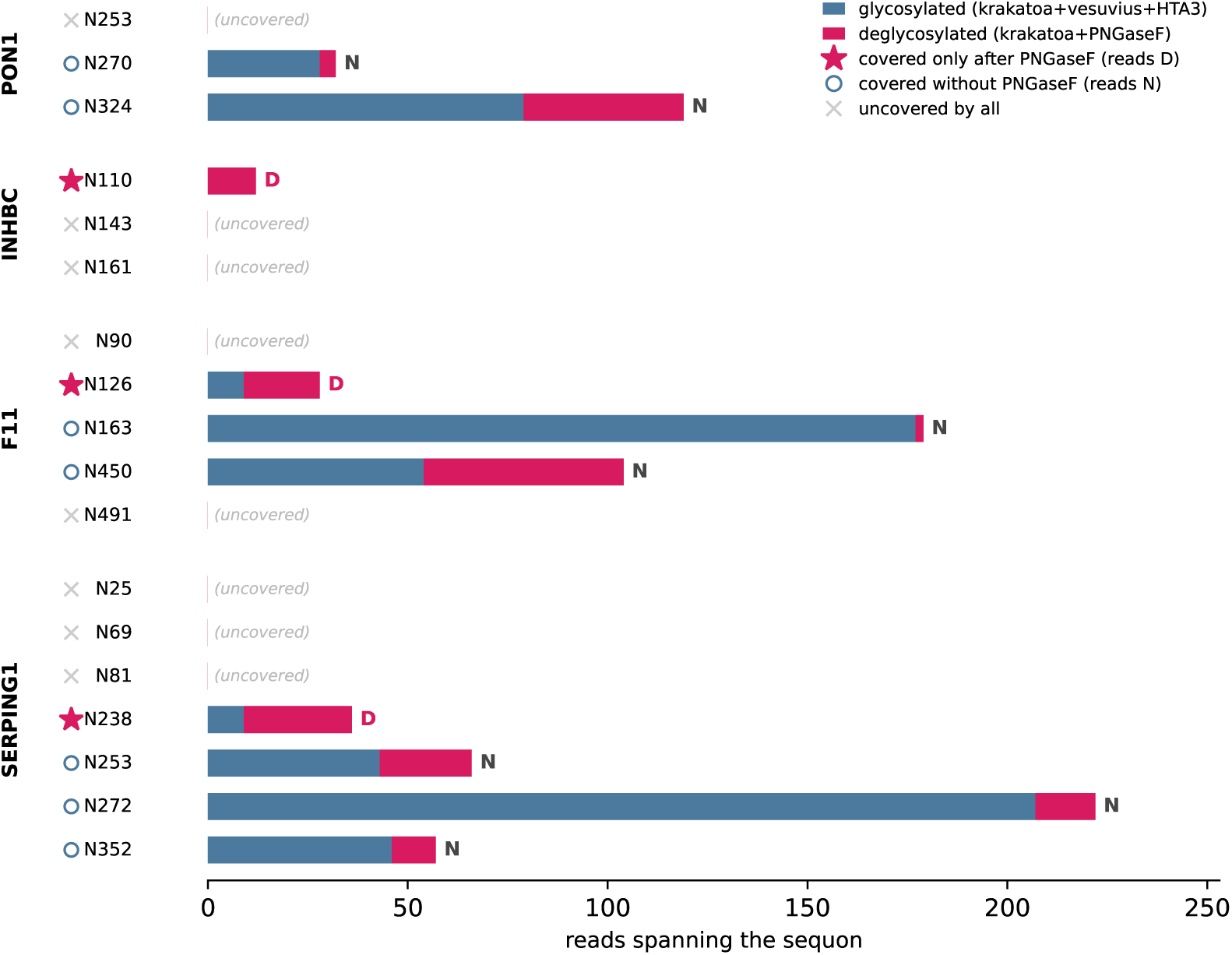
PNGaseF deglycosylation recovers the occupied N-glycosylation sites. For every annotated N-linked sequon in the five secretome proteins, the number of reads spanning it in the reference-free assembly, split between the three glycosylated proteases (krakatoa / vesuvius / HTA3, blue) and a krakatoa digest of a PNGaseF-deglycosylated aliquot of the same protein (magenta); the letter is the consensus residue called at the sequon asparagine. Markers flag whether a site is assembled only after the deglycosylated reads are added (*), already covered by the glycosylated digests alone (*◦*), or covered by neither (*×*). Adding PNGaseF raises the covered sequons from 7 to 10 of 18, and the recovery is specific to the occupied sites: every newly covered sequon reads aspartate (the PNGaseF deamidation product of a glycan-occupied asparagine), whereas the sites already covered read unmodified asparagine (their non-glycosylated peptidoforms). Eight sequons in low-yield regions remain covered by no digest.

The coverage tracks show *where* the reconstruction is right; read depth shows *why*. Each consensus column is one observation: the number of reads that voted into it, and whether the residue it calls matches the reference. Pooling these over all templates gives the relation directly (Figure 5), with the nine-protease antibody assembly and the three-protease panel on the same axis. Above about sixty votes the two datasets agree. At 62–91 reads both call 0.855 of columns correctly; at 137–203 they give 0.911 and 0.905, at 204–303 0.932 and 0.930. Column accuracy is therefore a function of depth, not of sample. Below sixty it falls steeply: in the panel a column carried by two reads is correct 22.7% of the time and one carried by six to eight 57.3%, against 93.4% at 92–136 and 99.1% above 304. Twenty votes give about 0.8, a hundred about 0.9. The assemblies differ in how much sequence reaches these depths: 522 of trastuzumab’s 1712 columns exceed 304 reads, whereas 470 of the panel’s 4368 fall below thirteen. Neither series has flattened at its deepest bin, so accuracy is limited here by attainable depth rather than by the consensus.

**Figure 5:**
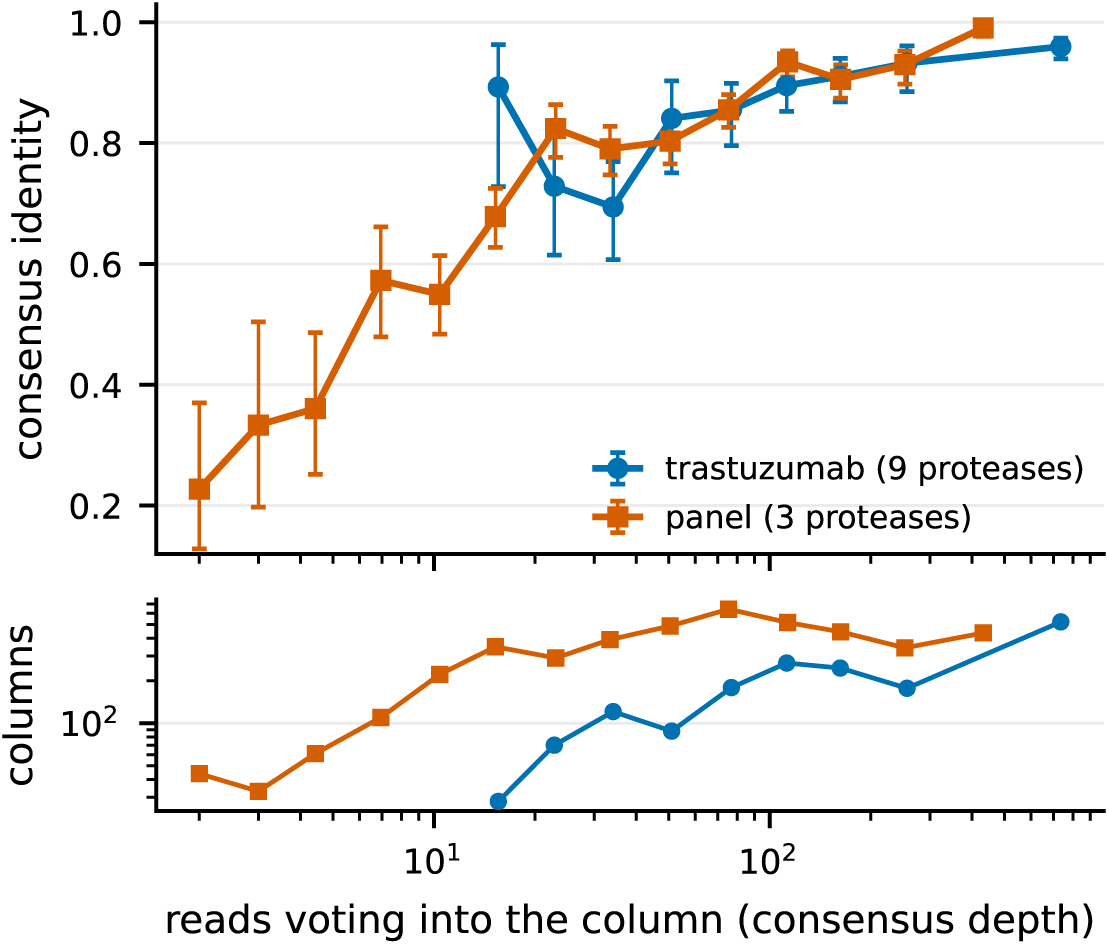
Consensus identity as a function of read depth. Every consensus column is one observation: the number of reads that voted into it, and whether the residue it calls matches the reference. Columns are pooled over all templates and binned geometrically by depth; each point sits at the geometric centre of its bin and the bars are Wilson 95% intervals. The two series are disjoint: the pooled nine-protease trastuzumab assembly, and the five non-antibody proteins of the three-protease panel (APOA4, PON1, F11, INHBC, SERPING1), the panel’s own trastuzumab digest being excluded so that no protein contributes to both. Column scoring follows the Table 3 convention, each template assigned to the reference it best windows onto and templates below 0.5 local identity dropped as background, except that here every template covering a position contributes rather than only the deepest, since the question is what a column of a given depth is worth. The lower panel gives the number of columns behind each point; bins holding fewer than ten columns are tabulated by the script but not plotted.

### 2.5 Comparison with prior antibody-sequencing methods

Antibody de novo sequencing has been driven by template-based assembly: de novo peptide reads are mapped onto a germline (IMGT) scaffold and a per-column consensus is read off, as in Stitch[15]. Systematic benchmarking of antibody de novo sequencing has begun recently[19]. We cite that effort as context rather than building on it: the comparison reported here is our own throughout, on our own data and with our own scoring.

The pipeline does not depend on a bespoke reader. borgonovo consumes any model that exposes a per-step softmax, so Casanovo[6, 7], ContraNovo[20] or InstaNovo[8] can each drive it unchanged, and we quantify what the reader contributes by substituting one. The results above use Pairwise Attention 1.1, an in-house model that on our six-protein panel exceeds Casanovo v5.2 by 0.084 fixed-window identity and 6 points of coverage; its fine-tuning corpus holds no spectrum from that panel and no trastuzumab-specific CDR sequence. The assembly gain is nonetheless separable from the reader: with either backend the consensus far exceeds the reads it is built from (0.454 to 0.792 with Casanovo, 0.470 to 0.876 with Pairwise Attention 1.1; Supporting Information), and on the nine-protease trastuzumab data, where the protease redundancy already saturates the reconstruction, the two backends are not separated (Casanovo 0.8671 against Pair- wise Attention 1.1’s 0.9000 fixed-window; difference +0.0329, 95% confidence interval *−*0.0214 to +0.0952, underpowered rather than null; these are per-contig scores against a chain-deduplicated reference and so are not directly comparable with Table 1).

Template-based assembly buys its accuracy with the germline scaffold, which engineered constructs lack. On the reference-free axis the only prior assembler we are aware of is ALPS[13], and we ran it ourselves, on our own reads, under a protocol registered before any comparison number was computed (Methods; the protocol, every grid cell and all scoring code are deposited).

Measured head-to-head, the two reference-free assemblers differ less in peak accuracy than in how reachable that peak is (Table 4). Scored on identical reads with a third-party metric (local BLOSUM62 alignment; precision over the full length of every contig, so unaligned and chimeric residues are charged, and recall over the reference), a fully tuned ALPS is the more accurate assembler on the deep nine-protease trastuzumab data: F1 0.986 from five contigs against borgonovo’s 0.954, its best cell over 465 configurations against borgonovo’s over 50. It is, however, a configuration selected by scoring against the reference sequence being reconstructed, which is precisely the information a reference-free method does not have.

**Table 4:**
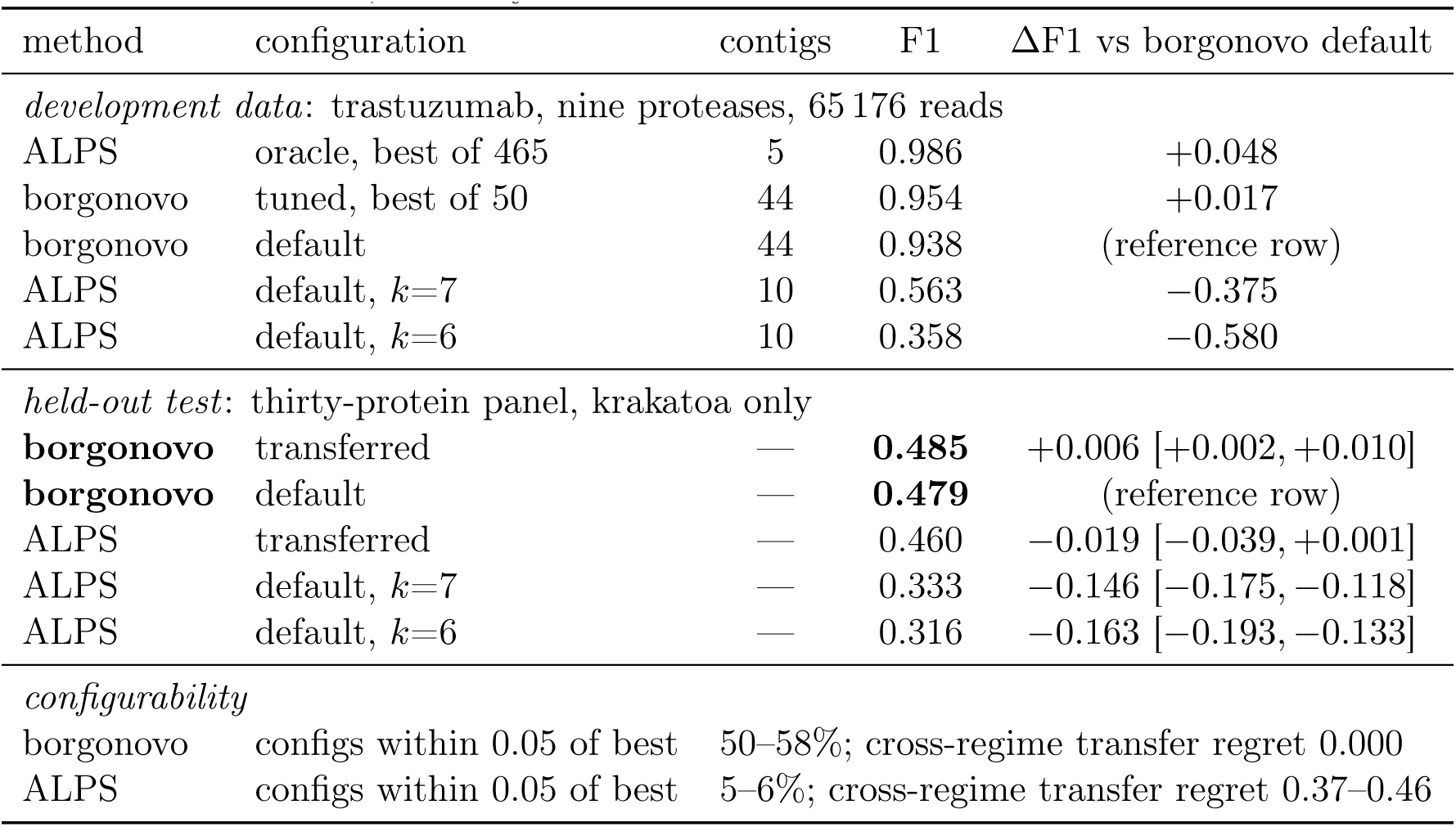
Measured head-to-head against ALPS[13]. Scoring is a third-party metric applied identically to both: local BLOSUM62 alignment, precision over the full length of all contigs (so unaligned, junk and chimeric residues are charged and no assignment threshold is used), recall over the reference, F1 their harmonic mean; identities fold I/L. Trastuzumab (nine proteases, HCD) is development data and was used to select both methods’ transferred configurations; the thirty-protein panel, digested with krakatoa alone, is held out, its sequences having played no part in configuring either method. The held-out block is therefore both a different set of proteins and a sparser, single-protease regime, and its absolute F1 values are correspondingly lower than the development block’s for both methods. *Oracle* marks a configuration chosen by scoring against the reference being reconstructed, which is not available to a reference-free user and is reported as an upper bound rather than an achievable operating point. *Transferred* configurations were selected on held-out Herceptin high-temperature digests by the same rule for both methods. ΔF1 is measured against the borgonovo default row of the same block, since the two blocks are different data and their anchors differ. *Default*means the configuration each method ships with: for borgonovo its published flags, and for ALPS its documented *c*=10 with the two k-mer lengths its manual recommends, since it names “6 or 7” without choosing between them and both are therefore shown. Confidence intervals are paired bootstraps over proteins, 10 000 resamples, and are therefore available only for the held-out block: the development block is a single protein, and resampling positions or contigs within one protein would understate uncertainty because neighbouring residues are carried by the same contigs. The development differences are point estimates with no interval and should not be read as significance tests. An interval lying wholly below zero means that row is resolvably worse than borgonovo at its default, and one spanning zero means the two are not separated by these data; the sign of the interval therefore carries the direction of the difference, and only the ALPS transferred row is unresolved.

We quantify that reachability three ways. First, ALPS ships no procedure for choosing its k-mer length: k is a required argument documented only as “6 or 7 is recommended”. Across our grid its F1 spans 0.03 to 0.99, only 5–6% of configurations lie within 0.05 of the best (borgonovo: 50–58%), and its documented default sits 33–40 points below its own optimum. Second, no reference-free proxy recovers the good setting: the natural heuristics are all inversely related to accuracy (Spearman *−*0.78 between true F1 and longest contig, *−*0.68 against total assembled residues), so maximising contig length selects the worst configuration in the grid, because small k builds long, chimeric and confidently wrong contigs. Third, a configuration tuned on one digest does not transfer across regimes: the trastuzumab optimum applied to the sparse single-protease high-temperature digests loses 0.37–0.46 F1, while borgonovo’s optimum is identical on both regimes, with a transfer regret of 0.000 and a spread of only 0.008 across all three datasets.

On thirty proteins whose sequences played no part in configuring either method, digested with krakatoa alone and so carrying far less redundancy than the nine-protease development data, borgonovo at its published default exceeds ALPS as distributed by 0.163 and 0.146 mean F1 (95% confidence intervals excluding zero; 29 of 30 and 30 of 30 proteins; sign test *p <* 10*^−^*^7^).

Given a configuration transferred from held-out development data, ALPS closes most of that gap and the two become statistically indistinguishable: borgonovo’s advantage is +0.019 with a confidence interval of *−*0.001 to +0.039, so by the pre-registered rule we downgrade the claim and assert no accuracy difference there. The pre-registered secondary metric splits the same way and we report the disagreement rather than resolve it: over the contigs clearing its assignment floor, transferred ALPS reaches higher per-residue identity than borgonovo (0.888 against 0.838 fixed, 0.917 against 0.902 fair) at 4.4 points lower coverage (36.9% against 41.2%). The consistent picture is that ALPS, correctly configured, produces fewer and cleaner contigs, while borgonovo covers more sequence at marginally lower per-residue identity.

The mechanism behind the brittleness is visible in ALPS’s implementation. Nodes are (*k−*1)- mers and edges are k-mers weighted by a normalised weighted geometric mean of per-residue confidence, accumulated additively over reads; contigs are then peeled by seeding at the heaviest node and extending greedily in both directions, consuming edges as the walk proceeds. There is no tip trimming, bubble popping, coverage thresholding or backtracking, so the walk commits irrevocably at every branch and a single spurious high-weight edge misroutes the remainder of a contig with nothing to detect or repair it.

The cross-regime failure, however, is driven less by *k* than by the second parameter. Nodes are exact (*k−*1)-mer strings, so a walk ends as soon as no read supplies an exact overlap, and mean contig length is therefore a dependent variable fixed by read depth rather than by the user: at matched *k* and prefilter we measure 113 residues per contig on the nine-protease data against 28 and 25 on the two single-protease digests. Since *c* caps the number of walks emitted, with the number of contigs equal to *c* in all 367 successful cells, the total sequence a run can output is approximately *c* times that depth-dependent length. Covering a 663-residue target thus needs *c ≈* 5 in the deep regime and *c ≈* 25 in the shallow one, and it is this budget, not *k*, that fails to transfer: carrying the deep configuration onto the sparse digests and correcting only *c* recovers 82.9% and 91.6% of the lost F1, whereas correcting only *k* recovers 0.0% and 5.5%. The asymmetry follows because under-budgeting removes reference positions one for one while surplus walks merely re-cover covered sequence, so a fortyfold excess on the deep data costs only 0.16 F1 at unchanged recall. *k* remains load-bearing through two secondary channels we also observe: below *k* = 7 repeat collapse in a walk with no visited-node guard produces grossly chimeric output, including a 5682-residue contig against a 663-residue reference, and above the optimum shorter contigs raise the required budget and strip the low-redundancy variable domains first, with variable-region recall falling from 0.978 at *k* = 9 to zero at *k* = 14 while the constant regions hold above 0.977. A practical consequence is that setting *c* generously is close to free and would remove most of the transfer penalty, so the deeper objection is not that ALPS cannot reach good assemblies but that neither of its two constants can be set from the data without knowing the answer. borgonovo has no *k* and no contig budget, and makes no irrevocable decisions: reads are aligned to a bootstrapped consensus and voted per column, so a misassigned read is outvoted rather than propagated, which is the structural reason its parameter surface is flat.

We do not claim to beat the template-based ceiling: on the same trastuzumab digests, of which we re-analyse only the HCD spectra, a template-plus-EThcD workflow reaches near-complete coverage and variable-region accuracy[21], which borgonovo (HCD only, no template) neither matches nor targets. Nothing in the method depends on the variable domain being known, the unknown-sequence case AbNovoBench identifies as the open problem for reference-free assembly, though every protein we report here has a reference sequence available, used only for scoring. Other database-independent routes exist, multi-protease contig scaffolding[16, 17] and mass-block overlap assembly with middle-down and intact-mass validation[18], and borgonovo differs from them in distilling a per-column consensus from the reads’ full softmax profiles using bottom-up data alone.

Three caveats bound the comparison. The identities fold I/L on both sides, so that axis is matched. Our figures are HCD-only, whereas template-based workflows are normally run with EThcD as well, which raises the ceiling available to them. And the comparison is run on our own trastuzumab, high-temperature and thirty-protein data; it is a same-data, same-metric benchmark against ALPS and nothing more. Trastuzumab additionally served to select both methods’ transferred configurations and is reported as development data, with the thirty-protein panel as the held-out test.

## 3 Discussion

The reference-free reconstruction is demonstrated on an antibody and on a panel of non-antibody proteins, but the consensus-prior re-decoding gain is so far reported only on one antibody protease’s spectra; running the second pass on the protein panel and an aggregate multi-protease evaluation are the natural next steps. On the broader panel the per-residue accuracy transfers intact, but the reconstructed extent is set by per-protein peptide yield: poorly expressed or heavily glycosylated proteins yield too few sequenceable peptides for any reference-free method, so coverage, not consensus accuracy, becomes the limiting axis for general application, addressed by deeper or more diverse digestion rather than by the assembler. A concrete instance is the trastuzumab heavy chain (Fig. 3, bottom left): with three proteases its constant region is cut only sparsely, into long and highly charged peptides, and 109 of its 450 residues go uncovered, in stretches of up to 52 residues. Searching the same spectra against the known sequence places a confident (*q <* 0.01) peptide over 95% of the uncovered residues, so the region is spectrally sampled; the limitation is in the de novo reading, not the acquisition. That figure is a property of the spectra rather than of any one backend: it is unchanged, to the decimal, between the two de novo models we compared, even though they leave slightly different regions uncovered. Of those sparsely-read spectra, 40% (mean peptide length 31 residues, mean charge 5+) yield no Casanovo read at all, and the remainder are decoded at only *∼*0.58 residue identity, frame-shifted or scrambled between anchored termini. With no clean, consistent reads to seed a template or vote into one, borgonovo leaves the tail uncovered rather than inventing sequence. Nine proteases recover the same region completely (Fig. 2), confirming that the remedy is deeper or more diverse digestion (or a de novo backend better at long, high-charge peptides) and not a change to the assembler.

Highly charged precursors are, on this evidence, the largest remaining source of unexploited signal. Just under a fifth of the panel’s spectra (19.8%) carry a precursor charge above 4+, and every result reported here gates them out. That gate is empirical rather than arbitrary: admitting charge 5–10 reads was tested on five de novo backends, two of them explicitly fine-tuned on high-charge spectra, and on every one of them it lowered mean per-residue identity (by 0.007 to 0.024 fixed-window) while raising mean coverage (by 0.2 to 5.7 points). The gate therefore buys identity at the cost of coverage rather than winning on both axes, and specialising the backend does not resolve the trade: the model fine-tuned most heavily on high-charge data loses the most identity (0.024) and gains the least coverage (0.2 points). The damage is done when such reads found or extend a consensus template, not when they vote into one. Down-weighting their votes is inert, seven weighting schemes moving every panel metric by less than 0.005, whereas barring the same reads from the scaffold at full vote weight recovers the whole difference; discarding them altogether is worse on both axes than letting them vote but not build. The high-charge fraction is therefore unexploited signal (these spectra are acquired, and a database search identifies peptides among them) that current de novo backends cannot yet read well enough to help a consensus. Closing that gap, whether by training on more and better-curated high-charge data or by architectures suited to long peptides, would raise coverage in exactly the sparsely- cut regions (such as the Fc tail above) where redundancy is currently thinnest. It is the most consequential improvement available to the pipeline, and it falls principally on the backend rather than on the assembler: controlling which reads may found a template limits the damage they do, but only a backend that reads these spectra can turn them into coverage.

The residual error is correlated across reads and therefore beyond consensus: it reflects the underlying de novo model, and would benefit from a stronger or differently-biased backend (borgonovo is backend-agnostic by design). Substituting Casanovo v5.2 for the Pairwise Attention 1.1 backend used throughout, the assembler and all of its settings unchanged, lowers the six-protein panel’s mean length-weighted fixed-window identity from 0.8761 to 0.7921 and its coverage from 82.0% to 76.0%: the backend accounts for +0.0840 of the identity reported above (95% confidence interval +0.0544 to +0.1129, stratified contig bootstrap). Overlap with the Pairwise Attention 1.1 fine-tuning corpus does not account for that margin. None of the panel’s own spectra entered the corpus, which holds no run from this acquisition; what it does hold are peptides of four of the six proteins observed in unrelated experiments (APOA4 in 275 spectra, F11 in 120, SERPING1 in 42 and PON1 in 28, counting peptides of at least nine residues) and none of inhibin *β*C. Trastuzumab matches 3963 spectra, but every one is human immunoglobulin constant-domain or germline-framework sequence, none covers a trastuzumab-specific CDR, and trastuzumab is absent from the database against which those spectra were identified, so no trastuzumab-specific peptide could have been labelled. Split on that criterion the mean gain is +0.068 over the four proteins represented and +0.115 over the two that are not. With only two proteins in the latter group that split is weak on its own, and the substantive control is that the benchmark spectra themselves are absent from the corpus. Sequence-level overlap of this kind is in any case hard to avoid and is not specific to one backend: any de novo model fine-tuned on human proteins will have seen peptides from abundant plasma proteins, and the same holds for the Casanovo checkpoints used here. What a reference-free benchmark requires is that the evaluated *spectra* were not used in training, not that the protein sequences were unseen. On the nine-protease trastuzumab data, scored per contig against a chain-deduplicated reference and so not directly comparable with Table 1, the same substitution lowers fixed-window identity from 0.9000 to 0.8671, but the contig bootstrap does not resolve that difference (95% confidence interval *−*0.0214 to +0.0952). Since that interval contains the panel effect, the single-protein comparison is underpowered rather than negative, and at the 635 reference positions both back- ends cover they agree closely (0.9701 and 0.9811): with nine proteases the redundancy already saturates the reconstruction, leaving little for the backend to add. A concrete, already-actionable instance is cysteine. On the free-thiol panel it is the worst-reconstructed residue (called correctly at only *∼*0.39 of covered positions, against *∼*0.79 for non-cysteine), and because its +57 Da mass mismatch misframes the neighbouring backbone the depression extends outward, recovering to baseline only *∼*4 residues away. This is not a consensus or tokenization failure but a training-data one: the backend has an unmodified-cysteine token yet never fires it, having seen almost only carbamidomethylated cysteine in training. The cysteine positions are only *∼*2.5% of panel residues, so the ceiling is modest (correctly calling every cysteine would raise panel fixed-window identity by *∼*0.01, or *∼*0.02 once the neighbour recovery is included), but it is a clean, self-contained gain that a backend trained to represent free cysteine would capture without any change to the assembler. Both halves of that diagnosis can now be measured. Over the panel runs Casanovo v5.2 decodes 1 832 077 residues and fires its bare-cysteine token exactly zero times, emitting the carbamidomethylated form 20 672 times (1.128%) instead, so on a non-alkylated sample every cysteine it calls carries a spurious +57 Da. Pairwise Attention 1.1, whose training labels were built with an explicit bare-cysteine mode and no fixed carbamidomethyl modification, fires the bare token 22 195 times (0.678%) against 11 592 (0.354%) for the modified form. Scoring the resulting consensus per reference position (a vote over the contigs assigned to each protein, so not on the scale of the *∼*0.39 quoted above), it calls 91 of 108 covered cysteines correctly, 0.843, against 60 of 85, 0.706, for Casanovo, narrowing the deficit relative to non-cysteine positions from 0.214 to 0.098. The predicted gain is therefore real, and it is obtained by representing free cysteine in the training labels.

That cysteine case illustrates a limitation of the backends rather than of the assembler, and it invites the question of whether a non-neural de novo sequencer could be used instead. Established tools such as PEAKS[22] are considerably more flexible than current deep-learning models about fragmentation modality, protease and variable modifications, which is where those models are weakest, and they report a per-residue local confidence. Nothing in the assembler requires a neural backend or HCD spectra: recruitment and voting need only a residue string with a per-residue confidence, which is what such a tool emits. We can bound what that would cost. Rebuilding every profile as the information-equivalent of a residue string plus one confidence per residue, with the argmax residue given that confidence and the remainder spread uniformly over the other residue tokens, and re-running the assembler unchanged, gives 0.939 against 0.938 on the full distributions (Methods). The assembly stage therefore does not exploit the full per-step softmax at all, and would be expected to work on confidence-annotated reads from any source, including a different fragmentation modality. What does require the distributions is the second-pass consensus prior, which blends the column posterior into the decoder and so needs a backend whose per-step probabilities can be intervened on. The natural division is thus a flexible sequencer for the reads and a neural backend only where re-decoding is wanted, and testing that division on PEAKS reads and on EThcD data is the most direct extension of this work.

Extending from a single mAb to low-complexity mixtures (2–4 antibodies) is the principal methodological challenge that remains: the present single-consensus, “do-not-fork” policy must give way to per-column allele calling that preserves genuine clonal branches as distinct templates. This is a recognized hard problem even for template-based assembly, where reconstructing the CDRH3 from dispersed polyclonal mixtures still requires manual curation of candidate sequences;[23] doing it reference-free is harder still. Germline/IMGT databases, deliberately unused here, could be layered on for absolute numbering and to rescue low-coverage stretches without compromising the reference-free core.

Redundant, overlapping de novo reads can be assembled reference-free into accurate protein consensus sequences. By bootstrapping a clean template from mass-closure-consistent reads and recruiting the remainder by substitution-tolerant voting, borgonovo reconstructs the trastuzumab Fab including its CDRs at 0.88–0.89 fixed-window residue identity (*∼*0.93 once local indels are accommodated) without any reference, and that consensus, fed back as a per-position prior, measurably improves the decoding of individual spectra.

## 4 Methods

### 4.1 Data

We used publicly available HCD MS/MS data for trastuzumab digested separately by nine proteases of differing specificity (aLP, AspN, chymotrypsin, elastase, GluC, LysC, LysN, thermolysin and trypsin), downloaded from ProteomeXchange under accession PXD023419.[21] Pooling the digests yields *∼*65,000 spectra whose peptides overlap extensively along the heavy (HC) and light (LC) chains. The trastuzumab HC/LC sequences are used only to evaluate the reference-free output; they are never used during profiling, assembly or re-decoding.

To test whether the approach generalizes beyond antibodies, we additionally analyzed a focused panel of five recombinant human plasma/secretome proteins together with trastuzumab itself, each digested separately by three broadly specific HTA proteases[3, 4] (krakatoa, vesuvius and HTA3) in triplicate and acquired by data-dependent HCD. The three replicate digests of each protein–protease pair were pooled, and each protein assembled both per protease and across all three; as before, each protein’s canonical UniProt sequence (where available) is used only for scoring, never during profiling or assembly. A broader 30-protein panel under a single protease, the one-protease arm of the same series, is described in the Supporting Information; four of its proteins are the secretome proteins above, so the two panels are not independent.

### 4.2 Panel digestion and LC–MS/MS acquisition

The five recombinant secretome proteins (PON1, INHBC, APOA4, F11 and SERPING1) were obtained from the human secretome resource of Tegel et al.[24], in which full-length secreted proteins are produced in Chinese hamster ovary (CHO) cells. Expression in a mammalian host means these proteins carry native-type N-linked glycans, which is what makes the occupied sequons systematically invisible to a bare-sequence reconstruction (Figure 4). Trastuzumab was obtained commercially (Roche). The three broadly specific HTA proteases, krakatoa, vesuvius and HTA3 (CinderBio), digest the proteins under the hyperthermoacidic conditions that define the class[3, 4]. HTA3 (CinderBio inventory number CB-23727) is a protease product still in development and not yet released commercially.

Each protein was diluted to 2 pmol/*µ*L in 60 mM citrate–phosphate buffer at pH 3 and digested in triplicate in a 96-well plate. Each 20 *µ*L reaction contained 20 pmol protein, 5 U of HTA protease and 5 mM tris(2-carboxyethyl)phosphine (TCEP) as reductant in the same buffer; vesuvius was added at three times the amount used for krakatoa and HTA3. Following the HTA- protease protocol the cysteines were not alkylated: at the pH 2–4 of the digest the reduced thiols are protonated rather than present as the thiolate required for thiol–disulfide exchange, so the disulfides do not re-form and no iodoacetamide or chloroacetamide capping step is required.

TCEP was chosen over dithiothreitol for the same reason, being an effective reductant at acidic pH. Digestion was initiated by transferring the plate to a thermoblock preheated to 80 *^◦^*C, run for 15 min at 80 *^◦^*C with orbital mixing at 600 rpm, and quenched by placing the plate on dry ice.

Because no alkylation was performed the panel cysteines are unmodified free thiols. The backend de novo model does carry an unmodified-cysteine token at the free-thiol mass, but, having been trained almost entirely on carbamidomethylated data, it effectively never emits it (over one Cys-rich panel run the token is the argmax in 0 of 53,660 decoding steps), defaulting instead to the +57.02 Da carbamidomethyl form. On these free-thiol samples every cysteine therefore carries a constant model–sample mass discrepancy, which makes cysteine the least accurately reconstructed residue and, because the misplaced +57 Da throws the local fragment-mass ladder out of register, also depresses the accuracy of its immediate neighbours (see Discussion). In a separate repeat experiment, krakatoa was used as the sole protease and the proteins were deglycosylated with PNGase F (Fisher Scientific, A39245) prior to digestion, cleaving the N-linked glycans so that the otherwise occluded sequons become accessible to the bare-sequence reconstruction.

Digests were analysed by nanoflow reversed-phase LC on an UltiMate 3000 RSLCnano system coupled through an EASY-Spray source to a Thermo Scientific Q Exactive HF Orbitrap mass spectrometer operated at 2 kV. 4 *µ*L of each sample was loaded onto a C18 Acclaim PepMap 100 trap column (75 *µ*m *×* 2 cm, 3 *µ*m, 100 Å) at 7 *µ*L/min and separated on an ES902 EASY-Spray PepMap RSLC C18 column (75 *µ*m *×* 25 cm, 2 *µ*m, 100 Å) at 0.7 *µ*L/min, both held at 60 *^◦^*C.

Solvent A was 3% acetonitrile / 0.1% formic acid in water and solvent B was 4.9% water / 0.1% formic acid in acetonitrile; the gradient was held at 1% B for 3 min then ramped to 32% B by 13 min, followed by a high-organic wash, with acquisition active over the first 17 min of the run. The instrument ran a data-dependent Top10 method in positive-ion mode. MS1 survey scans covered 300–1500 *m/z* at a resolution of 60,000 (AGC target 2 *×* 10^5^, maximum injection time 105 ms). Precursors below charge 2+ were excluded; selected precursors were isolated in a 2 *m/z* window and fragmented by higher-energy collisional dissociation at a normalised collision energy of 26, and MS2 spectra were acquired at a resolution of 30,000 (AGC target 1 *×* 10^5^, maximum injection time 105 ms) with dynamic exclusion for 20 s.

The panel comprises 54 runs: six proteins *×* three proteases *×* three replicate injections, acquired in a single batch and laid out one protein per plate row. The runs yielded 160,579 MS/MS spectra in total (median 2,887 per run). Precursor charge states were 2+ (39.0%), 3+ (29.9%) and 4+ (11.4%), with 19.8% of spectra above 4+, a substantial fraction that the de novo model does not attempt, and one source of the coverage ceiling discussed below. Only 4.4% of duty cycles reached the Top10 precursor cap, so acquisition was precursor- rather than method- limited, as expected for a purified single protein. Vendor raw files were converted to mzML with ThermoRawFileParser 1.4.5 with centroiding and no filtering, recalibration or other processing. All data generated and re-analysed here are deposited or cited under Data availability below.

### 4.3 First-pass profiling

Each spectrum is decoded greedily by a pretrained de novo model, the decoding backend, which nothing downstream is specific to. Rather than committing to a single sequence, we record at every decoding step the full softmax distribution over the residue vocabulary (the per-step *profile*). Profiles are variable length and are stored together with the precursor mass, charge and *m/z*, the model vocabulary and the order in which the decoder emits residues. Models that decode C*→*N, as Casanovo’s reverse tokenizer does,[6, 7] are re-oriented to N*→*C on loading, so assembly sees a single convention whichever backend produced the reads.

### 4.4 Reference-free template-bootstrap assembly

Let each read be the base-residue sequence of a profile’s per-step argmax (modifications stripped; isoleucine folded onto leucine, which is mass-identical). Assembly proceeds in three steps (Figure 1).

#### Scaffotd

We first select the cleanest reads by *mass closure*: a complete, correct decode’s summed residue masses plus a terminal group equals the precursor neutral mass. Reads passing closure within a ppm tolerance (here the densest mode of precursor *−* ladder) are the cleanest (*∼*74% residue accuracy on these data). Taking the highest-confidence such reads, we grow consensus templates progressively: each read is aligned (ungapped, substitution-tolerant, endoverhang-permitting) to the existing templates via a *k*-mer candidate prefilter; if its best identity clears a threshold it votes in, otherwise it seeds a new template. After a few reads have voted, a template’s consensus is cleaner than any individual read, so subsequent alignment is against a low-noise target.

#### Recruit

We then align *all* reads (not only the clean ones) to the fixed scaffold and confidence- vote into existing columns only, with no end-extension, so a noisy read cannot inflate a template. Reads spanning a framework*→*CDR junction anchor on the conserved framework and deposit votes across the hypervariable columns. Recruitment is iterated: the consensus is re-derived from the votes and all reads are re-aligned to the improved consensus, with the clean scaffold votes retained as an additive base. The recruit alignment can optionally be made *isobaric-tolerant* : beyond substitutions and short indels it credits an equal-mass block (a read’s GA against the template’s mass-identical Q, say) so that a read which tokenizes a residue differently from the template stays in frame and abstains at the ambiguous column instead of mis-voting it (the placement counterpart of the mass-aware scoring below).

#### Consensus and prior

Each template column accumulates the *summed full-vocabulary soft- max* of the reads aligned to it. Its argmax is the consensus residue; its normalization is a probability distribution over the next residue that doubles as a decoding prior. For each read we record its placement (template and column span) for the second pass.

#### Operating point

Every assembly reported here uses the same settings: a 50 ppm mass- closure tolerance for selecting the scaffold reads, the ungapped recruit aligner (the isobaric- tolerant variant is evaluated once, above, but is not the default), and columns whose read depth falls below 0.4 of their template’s median nonzero depth dropped at read-out (–closure-ppm 50 –prune-frac 0.4). Precursors above charge 4+ are gated out at profiling. No parameter is tuned per protein or per dataset.

### 4.5 Consensus-biased re-decoding

In a second pass, each spectrum is re-decoded greedily; at decoding step *s* the model softmax *m_s_* is linearly blended with the consensus column distribution *c_s_* at the spectrum’s recorded placement,

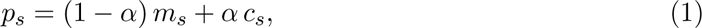

renormalized, with *α ∈* [0, 1] (default 0.5). The column is indexed by the decode step anchored at the read’s terminus, so the prior tracks the decode even if the re-decoded length differs slightly. Spectra with no placement decode normally (*α* effectively 0).

### 4.6 Mass-aware accuracy

Residue identity against a reference underestimates correctness for de novo MS in two ways: a single insertion/deletion misframes the rest of a contig under a fixed-length comparison, and distinct tokenizations of the same mass (e.g. Q vs GA, GG vs N) are physically indistinguishable from the spectrum. We therefore also report a *mass-aware* accuracy from a banded fitting alignment of contig to reference in which (i) a diagonal match/substitution step re-anchors the frame after a real error without a gap, (ii) an equal-mass block of up to *k* residues a side is credited as correct (capturing isobaric tokenizations), and (iii) the band bounds drift so the score cannot select a high-scoring subsequence. The difference between mass-aware and exact identity is the *isobaric-ambiguity share*.

That mass coincidence between isobaric residue combinations is a real source of de novo error has a long history in the field. Taylor and Johnson introduced a modified substitution matrix that treats single-residue isobars as equivalent when searching de novo outputs against a protein database.[25] Searle et al. implemented this as an explicit mass-based alignment (OpenSea), in which both de novo sequence and database sequence are converted to residue-mass series and aligned by comparing mass values, allowing blocks of residues with equal total mass to match.[26] Han et al. (SPIDER) and Zhang et al. (PEAKS DB) developed the same mass-block equivalence further for database-search identification.[27, 28] All of these approaches require a known protein database as the alignment target and work on top-1 or low-rank de novo sequences rather than full softmax distributions. Schulte and Snijder brought the mass-coincidence insight into the reference-free, read-*placement* setting by replacing sequence alignment with a mass-based alignment in Stitch, so that reads are placed without penalizing physically indistinguishable tokenizations.[23] We use the equal-mass move in both senses. Its primary use here is as a *scoring* operation, which partitions the apparent error of a finished contig into the isobaric- ambiguity share and the residual, genuine error; as reported below, on these data that partition is informative: mass coincidence accounts for only *∼*1% of the apparent error. The same move also serves *placement*, in the spirit of Schulte and Snijder: it is what makes the recruit alignment isobaric-tolerant (above), which modestly improves the reference-free reconstruction (below) at additional compute cost.

### 4.7 Per-contig scoring of the protein panel

Unlike the purified antibody, each panel sample also contains host-cell background, expression-tag remnants (a TEV cleavage motif was frequently observed) and low-level carry-over from neighbouring wells, so a contig need not belong to the intended protein. We therefore score *per contig*: each consensus contig is assigned to its best-matching reference among the panel proteins together with a common-contaminant set (cRAP), expression-tag motifs being flagged separately, and a protein’s coverage and identity are computed only over the contigs assigned to it. This separates the assembler’s per-residue accuracy from sample purity: a contig that correctly reconstructs a carried-over neighbour is credited to that neighbour rather than counted as an error against the intended protein. Coverage is the fraction of a reference spanned by its assigned contigs at *≥*50% local identity. Where a reference supplies the same chain more than once in near-identical form, as for an antibody heavy chain given with and without its C-terminal lysine, the variants are collapsed onto their longest representative before scoring: in a deep assembly both attract contigs and both would otherwise enter the coverage denominator, halving the reported coverage (on nine-protease trastuzumab, 96.4% against 60.5%). The *≥*50% floor is a deliberately permissive cut on assignment, not on per-residue quality (the identity reported per protein is the full residue accuracy of whatever clears it); it discards only contigs with no credible reference match: host-cell background, expression-tag fragments and chimeric mis-assemblies that align to no single protein. Of the assembled contigs, the great majority are assigned; the unassigned remainder do not enter any protein’s coverage denominator, so the panel identities are not inflated by silently dropping poorly-reconstructed contigs.

To mark, independently of the de novo backend, how much of each protein is in principle sequenceable from the acquired spectra (the “achievable” track beneath each coverage bar, Figure 3, and the uncovered-tail analysis in the Discussion), we searched the same spectra against each protein’s known sequence with a database engine (Tide/Percolator, Crux 4.3.2[29]), reporting peptides at a 1% peptide-level *q*-value. The search was non-specific (no enzyme assumed, matching the broadly specific digest) with a 15 ppm precursor tolerance, and carbamidomethylation of cysteine was specified as a *variable*, not fixed, modification: the HTA protocol reduces with TCEP and does not alkylate, so cysteine is a free thiol, and a fixed +57 Da carbamidomethyl assumption makes every cysteine-containing peptide 57 Da too heavy and therefore unidentifiable. With the variable modification the number of confidently identified cysteine-containing peptides across the panel rises roughly tenfold, and 95.8% of the cysteines in identified peptides are found unmodified, confirming the free-thiol chemistry directly from the data. The effect scales with cysteine content: the cysteine-free apolipoprotein A-IV is unchanged, whereas the achievable coverage of the cysteine-rich factor XI rises from 76% to 98%, so a fixed-modification search would have understated the sequenceable ceiling most severely where disulfide-bonded cysteine is densest.

### 4.8 Head-to-head comparison against ALPS

#### Pre-registration

The protocol was written, committed and tagged before any borgonovo- versus-ALPS number was computed (reproduce/protocols/alps_baseline.md, tags alps-protocol-v1, v1.1, v2-matched and panel30-configs-frozen). It fixes the read sets, the exporter, the grids, the selection rules, the primary metric, a gate that voids the comparison if ALPS is crippled, and an outcome table covering the case where ALPS wins. Two deviations are recorded there: exploratory ALPS runs made during feasibility scoping, whose numbers are not reported, and an extension of the ALPS grid beyond the originally registered axes after its optimum landed on two boundaries with F1 still rising. The extension can only help ALPS.

#### Software

ALPS.jar (md5 6e1062c08236723a9185f1c773976b84) was obtained from the Scientific Reports supplement of ALPS[13] (41598_2016_BFsrep31730_MOESM2_ESM.zip, CC BY 4.0) and run unmodified under OpenJDK 21.0.11. As an installation control it reproduces its own published human light-chain assembly byte for byte. The jar carries no software licence and the weighted de Bruijn algorithm is under patent US10309968B2, so it is fetched at run time by reproduce/scripts/fetch_alps.sh rather than redistributed, and is invoked as a binary rather than reimplemented.

#### Input bridge

reproduce/scripts/export_alps.py writes the positional CSV ALPS parses (identifier, peptide, space-separated per-residue confidence, intensity) from the same profile .npz files borgonovo consumes, so both assemblers see identical reads. Confidence is the per-step argmax posterior scaled to ALPS’s 0–100 column. Five encodings were swept; because the argmax of a softmax over 45 tokens is bounded below by 1*/*45 and never fell under 0.07, flooring at 5, at 1 or not at all gave identical results. As a positive control, in-silico perfect reads digested from the reference and passed through the identical bridge reconstruct both chains at 100% coverage, so the bridge is not the source of any observed limitation.

#### Metric

The primary metric imports nothing from borgonovo: Biopython PairwiseAligner, local mode, BLOSUM62, gap open *−*11, extend *−*1. Precision is summed aligned identities over the summed full length of all contigs, so junk, unaligned and chimeric residues are charged and no assignment threshold is used; recall is the union of reference positions matched by identity over reference length; F1 is their harmonic mean. borgonovo’s own panel_score is reported as a pre-registered secondary, with the number of proteins clearing its 0.5 assignment floor stated, because that floor removes contigs from both numerator and denominator asymmetrically between methods.

#### Configuration and the devetopment/test sptit

ALPS was swept over 465 configurations on trastuzumab and 100 on each high-temperature protease; borgonovo over 50 on each. Because ALPS provides no procedure for choosing *k*, and because selecting it against the reference is not available to a reference-free user, both methods’ reported panel configurations were selected on held-out Herceptin high-temperature development data by the same rule, best worst-case F1 across the two proteases (krakatoa and vesuvius), and then frozen. The thirty-protein panel, digested with krakatoa alone, is the test set; none of its sequences informed either configuration. Selecting on Herceptin HTA digests rather than on the nine-protease data was deliberate, since it matches the single broadly specific protease and the read depth of the test set: a configuration transferred instead from the deep nine-protease data is penalised by 0.37 to 0.46 F1 in this regime, which would have handicapped ALPS for a reason unrelated to its assembly algorithm. Oracle configurations are reported where stated as upper bounds only.

#### Statistics

Paired bootstrap over proteins, 10 000 resamples, with the same resampled index set applied to both arms so pairing is preserved, plus exact sign and Wilcoxon signed-rank tests. The protocol pre-committed that a confidence interval including zero downgrades the claim, which is what occurred for borgonovo’s default against a transferred ALPS.

#### Controts

Beyond the reproduction and perfect-read controls above, four pre-registered controls were run. A *constant-region gate* required tuned ALPS to reach *≥*0.90 recall on the constant domains or the comparison would have been declared crippled and discarded; it reached 0.979.

A *uniform-rescale* check confirmed that multiplying every confidence by an exact factor leaves ALPS’s output byte for byte unchanged, as its normalised weighted geometric mean predicts. A *depth titration* over 10–100% of reads, three seeds each, separates the two methods sharply:

ALPS at *k*=9 rises monotonically with depth (0.907 to 0.986) as its k-mer graph fills in, ALPS at *k*=7 is erratic with seed-to-seed standard deviations up to 0.10, whereas borgonovo varies by at most 0.010 between seeds and is nearly flat in depth. An *information-parity ablation* rebuilt each profile as the ALPS-equivalent posterior (argmax residue receives the confidence, the remainder spread uniformly over the other residue tokens) and re-ran borgonovo unchanged: it scores 0.939 against 0.938 on the full softmax, so its margin is not an artefact of consuming a richer read representation. One control returned a null result: permuting per-residue confidences within each read left ALPS’s F1 unchanged at 0.9856, indicating that at its selected operating point, where a mean-confidence prefilter has already retained only uniformly high-confidence reads, the confidence column itself carries little information and the assembly is driven by k-mer topology and read selection. Because the permutation preserves each read’s multiset of confidences, this bounds rather than excludes a role for the confidence weighting.

### 4.9 Implementation and availability

borgonovo is a Python package that adds the profiling, assembly and re-decoding logic externally to a de novo sequencer, which it treats as an interchangeable component: any model exposing a per-step softmax can drive the pipeline with no change to the assembler. The backend used throughout, and against which the results below are reported, is Pairwise Attention 1.1, a retrained model built on the mass-difference attention of Lapin et al.,[9] released with this work. It can be replaced by Casanovo[6, 7], which the package consumes as a library and offers as its default backend, and which we use as the substitution control quantifying how much of the reconstruction the backend supplies, or by another model such as InstaNovo[8].

## Acknowledgements

LK was supported by the Swedish Research Council (Vetenskapsrådet), grant VR-2024-05887 and Knut and Alice Wallenberg Foundation KAW 2022.0032. The training of de novo sequencing transformers was enabled by the Berzelius resource provided by the Knut and Alice Wallenberg Foundation at the National Supercomputer Centre.

## Author contributions

A.N. trained the Pairwise Attention 1.1 model. E.S. performed the sample preparation and LC–MS/MS acquisition and wrote the corresponding Methods. D.S. and J.S. contributed to the development of the assembly algorithm through discussion and to the writing of the manuscript.

F.E. contributed to the conception of the study, introduced the high-temperature protease chemistry on which the digestion protocol rests, and provided mass spectrometry resources. L.K. conceived the method, implemented the software and wrote the manuscript. F.E. and L.K. supervised the work. Claude Code (Anthropic) was used as an assistive tool, both in implementing the software and in drafting and revising the manuscript. All authors read and approved the final manuscript.

## Data availability

The mass spectrometry proteomics data generated here, comprising the six-protein three-protease panel, the PNGase F-treated krakatoa runs and the 30-protein single-protease panel, have been deposited to the ProteomeXchange Consortium via the PRIDE partner repository[30] with the dataset identifier PXD082222. That deposit also carries the components of the Pairwise Attention 1.1 fine-tuning corpus that were generated here, which share runs with the 30-protein panel, so the training overlap reported in the Supporting Information can be verified against the deposited files themselves. The remaining component of that corpus is the previously published plasma protein-corona dataset PXD007648[31], which we reuse without re-depositing. The trastuzumab data re-analysed here are the published nine-protease set of Peng et al.[21] The Pairwise Attention 1.1 model checkpoint is deposited in the SciLifeLab Data Repository[32].

## Code availability

borgonovo, including the analysis scripts used to produce the figures and tables reported here, is available at https://github.com/statisticalbiotechnology/borgonovo under the Apache- 2.0 licence.

## Supporting Information

### Reconstructed sequences

The Results section reports coverage and identity as aggregates. Below is the sequence borgonovo actually called, residue by residue, against the known reference, for apolipoprotein A-IV and for both trastuzumab chains. APOA4 is the more representative case: it is a three-protease HTA digest of the kind the method is aimed at, and the best-covered protein of the panel, whereas the trastuzumab reconstruction draws on nine conventional proteases. Scoring is the convention behind Figures 2 and 3 — each template is assigned to the reference it best windows onto and the deepest covering template calls each position — and no reference is used at any point in producing the sequence, only in colouring it here. I and L are folded throughout: they are isobaric, so a de novo reader cannot separate them and calling one is not an error.

Two things are worth reading off these listings rather than the percentages. First, the errors are not spread evenly: they cluster into a few short runs, and the long constant stretches are recovered exactly. Second, a run of apparent miscalls is often a single local indel rather than many wrong residues — at the APOA4 mature N-terminus the contig is placed one residue early, so thirteen positions read as wrong while the underlying sequence is right. This is precisely the misframing that the fixed-window metric cannot absorb and the fair (indel-absorbing) metric does, and it is why the two differ by *∼*0.08 for APOA4 (Table 3) against *∼*0.05 on trastuzumab.

**APOA4 (P06727).** 396 aa, 96.0% covered, 94.7% of covered residues correct.

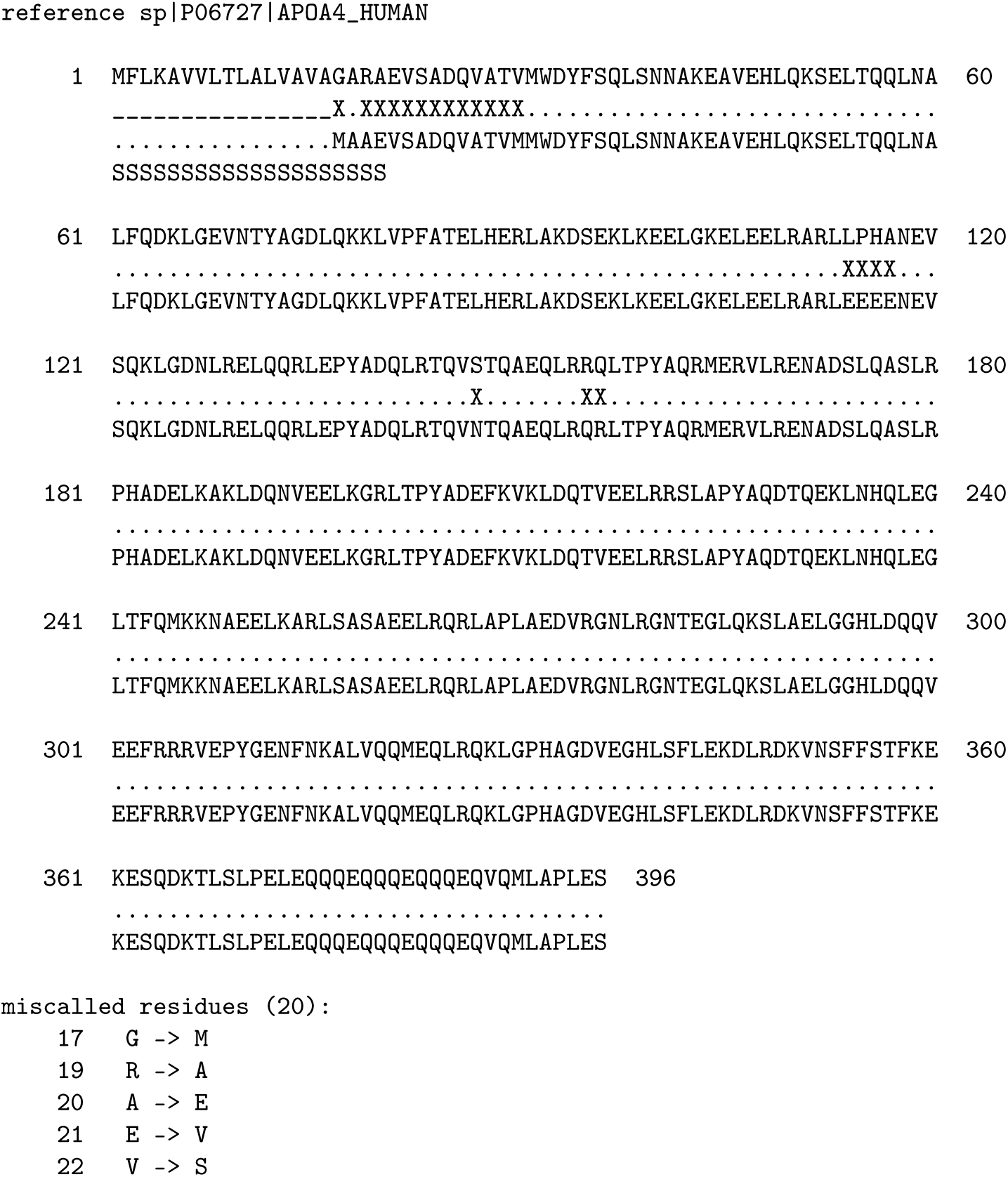
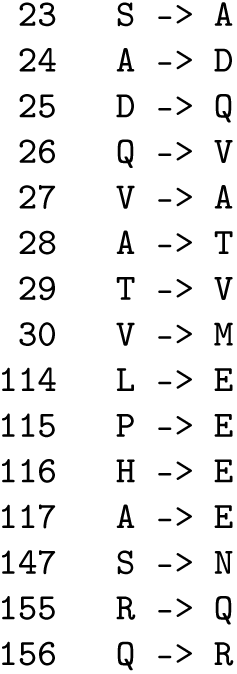

Trastuzumab heavy chain. 450 aa, 100.0% covered, 96.9% of covered residues correct.

reference trastuzumab_Herceptin_heavy_chain_plus_C_terminal_K_variant_450aa

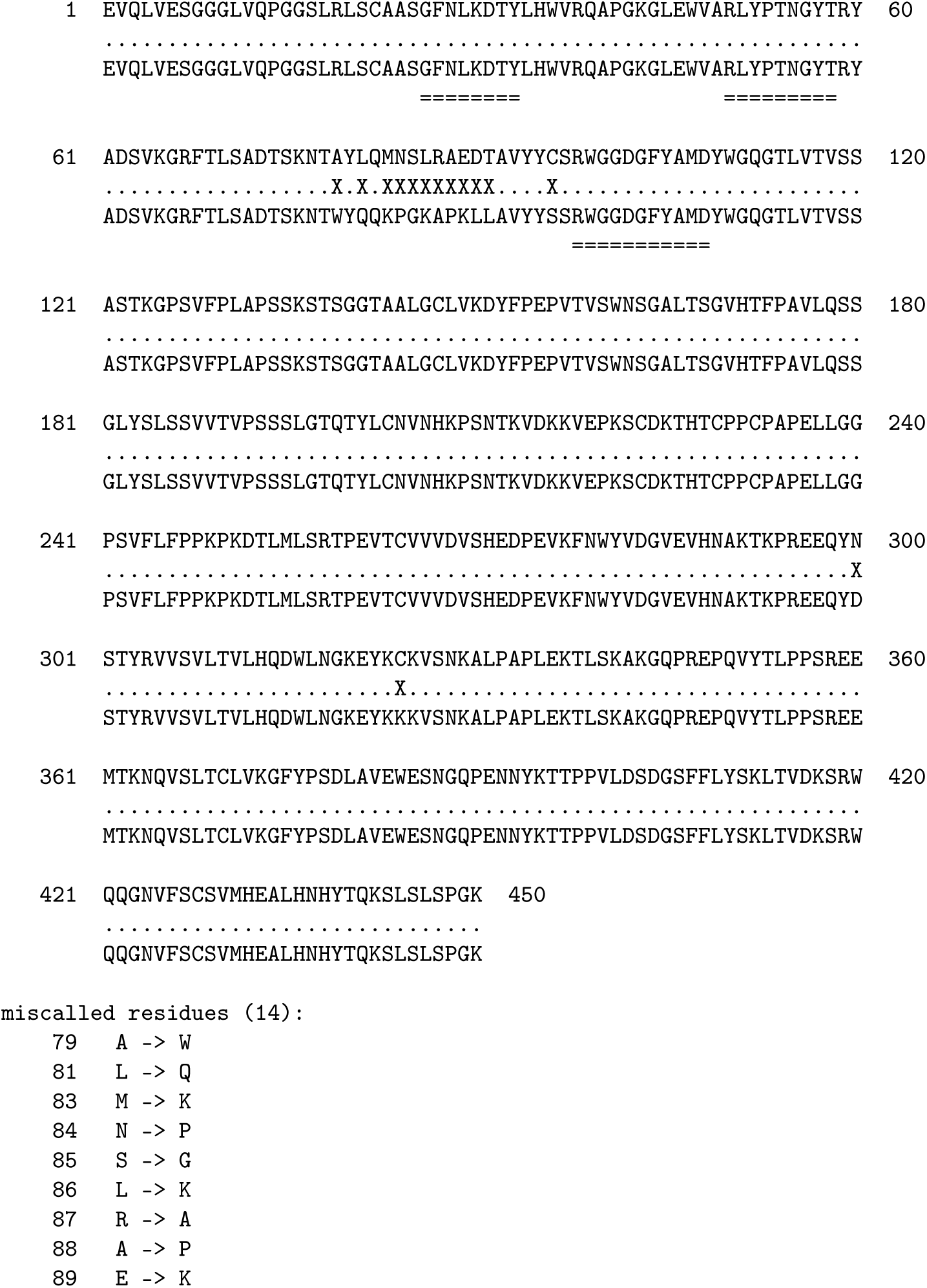
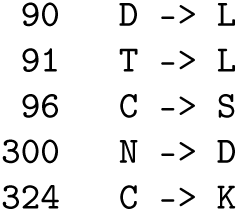

Trastuzumab light chain. 214 aa, 100.0% covered, 95.3% of covered residues correct.

reference trastuzumab_Herceptin_light_chain_WHO_RL65_214aa

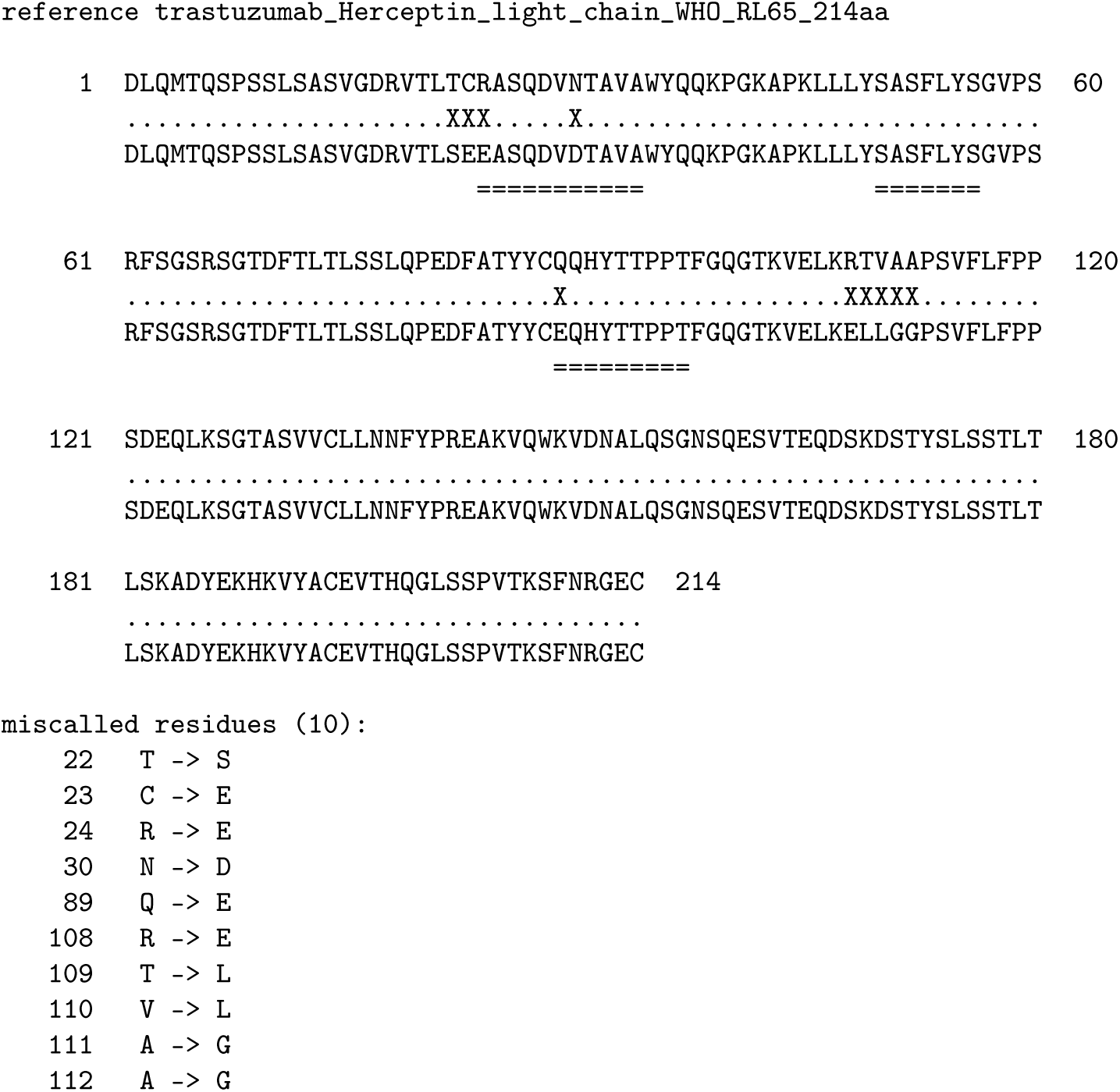

### Singte-protease breadth arm: 30 proteins

borgonovo was additionally applied to a panel of 30 recombinant human plasma/secretome proteins (128–1208 residues) under a single broadly specific digestion, with krakatoa alone; Table S1 lists the per-protein reconstruction. This is the one-protease arm of a protease-count series the main text otherwise samples only at three and nine, and since krakatoa is also one of the three proteases of the main panel the series is nested in its protease set: mean coverage is 39.5% with one protease, 82.3% with three (Table 3) and 100.0/100.0% on the nine-protease trastuzumab heavy and light chains (Table 1), while per-residue identity stays essentially flat throughout. Across this panel the identity is a mean 0.82 fixed-window, 0.89 fair and 0.92 mass-aware over the covered regions, matching the three- protease panel and trastuzumab over a *∼*10-fold range of protein length. The input Casanovo reads average only 0.47 fixed-window identity against their protein (*∼*8% are *≥*0.9 correct), so the consensus they vote into (0.82) is a *∼*1.7*×* per-residue improvement on the matched comparison (quoted like-for-like; the conservative floor is given below for the six-protein panel, where the comparison was repeated), while a naive overlap assembly of the same reads scores below the reads themselves (0.14–0.41). Coverage here (mean 39.5%, of a mean 52% reachable and 48% assemblable by any read) is bounded by per-protein peptide yield rather than by the assembler. That matched comparison does not depend on this panel. Repeating it on the six-protein panel, whose spectra enter no training corpus used here, reproduces it closely: its 115 174 Pairwise Attention 1.1 reads average 0.4699 fixed-window identity against their protein with 11.4% at least 0.9 correct, against a consensus of 0.8761, a 1.86*×* improvement (1.18*×* under a matched assignment floor, see below). Substituting Casanovo v5.2 lowers both sides and the multiplier with them (reads 0.4538, 8.4% at least 0.9 correct, consensus 0.7921, 1.75*×*; 1.09*×* matched), so the stronger backend does not merely hand the consensus better reads to inherit, it also leaves more for the consensus to add. The two sides are filtered differently, since a contig enters the consensus score only above the 0.5 assignment floor while every read is counted; imposing that floor on the reads as well is stricter than the assignment rule warrants here, each panel run being a single purified protein, so we quote the unfiltered 1.75*×* and 1.86*×* as the like-for-like figures and the matched 1.09*×* and 1.18*×* as a conservative floor. The consensus exceeds its own input reads on either convention.

Four of the 30 (APOA4, PON1, FA11 = F11 and IC1 = SERPING1) are also main-panel proteins, digested there by three proteases instead of one, so they give a matched within-protein contrast: coverage rises 79.5*→*90, 44.8*→*85, 24.6*→*76 and 52.8*→*68% respectively, a mean 50.4*→*79.8% (+29 points), while mean fair identity is unchanged (0.914 against 0.895).

Added proteases buy coverage; the consensus already supplies identity from a single digest. The 30 should therefore not be read as 30 independent proteins additional to the main panel.

#### Backend and provenance

This panel was profiled and assembled with Casanovo v5.2 throughout. No Pairwise Attention 1.1 results are reported on it: the backend comparison is confined to the six-protein panel, where the fine-tuning provenance of every protein is known. All 30 of these proteins contribute peptides to the Pairwise Attention 1.1 fine-tuning corpus, and the overlap is stronger than sequence alone: for 26 of them the corpus contains this panel’s *own spectra*, 25 038 of them drawn from 75 of the panel’s 86 runs, identified by database search and used as training labels. The four exceptions are exactly the proteins this panel shares with the six-protein benchmark (APOA4, PON1, FA11 = F11 and IC1 = SERPING1): they were held out of that conversion, so no spectrum of a benchmark protein entered the corpus by this route, and they appear only as peptides observed in unrelated experiments (275, 28, 120 and 42 spectra respectively). Because every figure reported for this panel is a Casanovo figure, none of them is affected by that overlap; the panel could not, however, be used to evaluate Pairwise Attention 1.1, which is why the backend comparison is confined to the six-protein panel, whose own spectra appear nowhere in the corpus.

---

**Table S1:** Per-protein reference-free reconstruction of the 30-protein single-protease panel, decoded with the Casanovo backend throughout. Coverage is the percentage of the UniProt reference spanned by the contigs assigned to that protein (*≥*50% local identity); fair identity is the indel-absorbing per-residue identity over those contigs (“—” where no contig was assigned). Biotinidase (BTD) yielded essentially no sequenceable peptides. Identity is computed over covered residues only, so for the least-covered proteins it rests on few residues (e.g. F13B’s 1.000 spans *∼*77 of 661): coverage and identity should be read together, not separately. APOA4, PON1, FA11 and IC1 also appe<u>ar in</u> Table 3 <u>under a three-protease digest</u>.

| protein | UniProt | len | cov.% | fair id. |
| --- | --- | --- | --- | --- |
| APOA4 | P06727 | 396 | 79.5 | 0.864 |
| APOM | O95445 | 188 | 55.9 | 0.876 |
| BTD | P43151 | 464 | 0.0 | — |
| C4BPB | P20851 | 252 | 19.4 | 0.962 |
| CLUS | P10909 | 449 | 48.8 | 0.874 |
| CO8B | P07358 | 591 | 11.5 | 0.857 |
| CPN2 | P22792 | 545 | 12.5 | 0.947 |
| CXCL7 | P02775 | 128 | 18.0 | 0.859 |
| F13B | P05160 | 661 | 11.6 | 1.000 |
| FA11 | P03951 | 625 | 24.6 | 0.879 |
| FBLN5 | Q9UBX5 | 448 | 22.1 | 0.913 |
| HEP2 | P05546 | 499 | 63.7 | 0.909 |
| HPT | P00738 | 406 | 38.7 | 0.907 |
| IC1 | P05155 | 500 | 52.8 | 0.930 |
| ICAM2 | P13598 | 275 | 48.7 | 0.868 |
| IL10 | P22301 | 178 | 60.1 | 0.882 |
| IL6 | P05231 | 212 | 78.3 | 0.874 |
| IPSP | P05154 | 406 | 4.2 | 0.656 |
| KAIN | P29622 | 427 | 75.2 | 0.898 |
| MSLN | Q13421 | 630 | 19.0 | 0.920 |
| NCHL1 | O00533 | 1208 | 35.2 | 0.907 |
| PCSK9 | Q8NBP7 | 692 | 44.5 | 0.873 |
| PON1 | P27169 | 355 | 44.8 | 0.984 |
| PRL | P01236 | 227 | 67.4 | 0.887 |
| Q9UNN8 | Q9UNN8 | 238 | 56.3 | 0.861 |
| QSOX1 | O00391 | 747 | 40.7 | 0.916 |
| TETN | P05452 | 202 | 43.1 | 0.914 |
| THBG | P05543 | 415 | 63.6 | 0.919 |
| THRB | P00734 | 622 | 12.2 | 0.885 |
| VASN | Q6EMK4 | 673 | 31.6 | 0.898 |

